# *Plasmodium falciparum* Myosin B is a slow motor optimized for force generation

**DOI:** 10.64898/2026.09.11.750570

**Authors:** James P. Robblee, Dihia Moussaoui, Carol S. Bookwalter, Daniel Auguin, Patricia M. Fagnant, Jill E. MacFarlane, Michael J. Previs, Anne Houdusse, Kathleen M. Trybus, Julien Robert-Paganin

## Abstract

Malaria is a disease caused by apicomplexan parasites of the genus *Plasmodium*. These organisms express two atypical class-XIV myosins: Myosin A (MyoA), a core component of the glideosome expressed throughout the entire lifecycle, and Myosin B (MyoB), which is restricted to invasive stages and localizes to the apical region. Here, we determine the crystal structure of *Plasmodium falciparum* MyoB (PfMyoB) in the Rigor state and identify its essential light chain as the same subunit bound to MyoA. Combining structural analysis, molecular dynamics simulations and in vitro kinetic and motility assays, we show that PfMyoB is a slow motor optimized for force production during invasion. The N-terminal extensions of PfMyoA and PfMyoB exert distinct effects on each motor’s mechanochemistry. Altogether, these findings reveal how Plasmodium myosins have evolved specialized functions during the complex parasite lifecycle and provide insight into developing multi-target inhibitors of erythrocytic invasion based on the PfMyoA inhibitor KNX-002.

## Introduction

Malaria, a blood-borne infectious disease caused by parasites of the genus *Plasmodium* remains a major global health burden, accounting for more than 600,000 deaths annually (WHO report, 2024). Parasite gliding motility and host red blood cell invasion rely on an actomyosin motor system, making parasite myosins attractive candidates for drug development. Myosins share a conserved architecture consisting of an N-terminal motor domain responsible for actin binding and ATP hydrolysis, a lever arm that binds calmodulin (CaM)-like light chains, and a variable C-terminal tail region that drives self-assembly and/or recruitment of binding partners and cargo^1,2^. The actin- and ATP-dependent cycle of myosin begins when the motor binds ATP, detaches from F-actin, and enters the **post-rigor (PR) state**. During the recovery stroke, ATP is hydrolyzed and the lever arm is primed, allowing the motor to reach the **pre-powerstroke (PPS) state**. The motor then rebinds actin and performs the powerstroke when phosphate (Pi) and Mg.ADP are sequentially released, resulting in the nucleotide-free **Rigor state**. The myosin is then ready to bind Mg.ATP, detach from actin and initiate a new cycle (reviewed by^2^)(**Supplementary Fig. 1a**). Variations in the rates and equilibria between these states, as well as unique structural features of different myosins, contribute to the diversity of motor functions.

Six myosins are expressed in *Plasmodium*^3,4^, including the two shortest unconventional class XIV myosins, myosin A (PfMyoA) and myosin B (PfMyoB). PfMyoA is part of the glideosome, a macromolecular complex that drives gliding motility and invasion of red blood cells^5^. PfMyoA binds two light chains, the essential light chain (PfELC)^6,7^ and the myosin tail interacting protein (MTIP), whose N-terminal region tethers the motor to the GAP complex and functionally acts as a “tail”^8^. PfMyoA is an exciting pharmaceutical target against malaria because it is essential for erythrocytic invasion in *P. falciparum*^9^. The therapeutic potential of PfMyoA has been further supported by the recent development of small-molecules capable of inhibiting its activity and blocking parasite pathogenesis: KNX-002 and KNX-115^10,11^.

PfMyoB is expressed in all invasive parasite stages of the parasite and localizes to the apical tip of the merozoite, in contrast to PfMyoA that spans the periphery of the parasite just beneath the plasma membrane^12,13^. Deletion of PfMyoB results in a delay in the initiation of parasite internalization, indicating that this motor contributes to invasion but is not essential^14^. A single light chain has been identified for PfMyoB, myosin light chain-B (MLC-B), which consists of a canonical calmodulin (CaM)-like domain and a long helical N-terminal extension^12^. MLC-B is essential and is expressed prior to MyoB, indicating that it likely has additional interacting partners besides PfMyoB^12,13^. *In vitro* studies have shown that PfMyoB binds actin filaments and has actin-activated ATPase activity^15^.

Recent studies of PfMyoA have revealed an atypical mechanism for tuning force production, in which a phosphorylatable serine (Ser19) within the N-terminal extension of the heavy chain binds to and stabilizes the Converter in the Rigor conformation. This interaction compensates for the reduced flexibility of the SH1-helix caused by a glycine to serine substitution at the “Fulcrum” hinge position^9^(**Supplementary Fig. 1b-d**). Ser19 phosphorylation directly modulates motor properties: when phosphorylated, the motor moves actin filaments at high speed, whereas when dephosphorylated, the motor is slower but produces more force^9^. Subsequent *in vivo* experiments have demonstrated that the phosphorylation state modulates parasite motility and force generation^16^. Despite their common classification as class XIV myosins, PfMyoB and PfMyoA are highly divergent, sharing only ∼35% sequence identity and displaying major differences in the sequence of their allosteric connectors and N-terminal extensions (**Supplementary Fig. 1d**). These differences suggest that the two motors may rely on distinct mechanisms of force production.

Here, we show that expressed PfMyoB binds two light chains: its unique MLC-B and the same PfELC bound by PfMyoA. By combining structural and functional analyses, we demonstrate that the force production mechanism of PfMyoB markedly differs from that of PfMyoA. Unlike PfMyoA, PfMyoB properties are unchanged by phosphorylation of the N-terminal heavy chain extension. PfMyoB is a constitutively slow myosin, tuned for ensemble force production and characterized by a slow nucleotide-binding rate and weak nucleotide affinity that depend on its N-terminal extension. This work reveals how two motors from the same class can diverge to fulfill distinct physiological functions, highlighting the diversity and adaptability of myosin motors. We also demonstrate that while KNX-002 exerts only modest inhibition of PfMyoB ATPase activity, the PfMyoB crystal structure provides a framework for the future design of KNX derivatives capable of targeting both PfMyoA and PfMyoB, the two motors involved in erythrocytic invasion.

### PfMyoB heavy chain binds MLC-B and PfELC light chain

MLC-B, the only characterized light chain for MyoB, binds to the more C-terminal IQ motif of the heavy chain^12^. The light chain that binds to the first IQ motif of the PfMyoB heavy chain was unknown despite attempts to identify it^15^. Similar to PfMyoA, the two IQ motifs of PfMyoB are non-canonical, suggesting that they bind unconventional light chains through distinct binding modes^17^(**Fig. 1a**). We identified the essential light chain that occupies the first IQ motif by co-expressing 10 potential light chains (**Supplementary Table 1**, Methods) with MLC-B and a tagged PfMyoB heavy chain in *Sf*9 cells and isolating the complex by affinity purification. Only PfELC, previously identified as the essential light chain of PfMyoA^6^, bound tightly enough to co-purify with the PfMyoB heavy chain.

When PfMyoB heavy chain is co-expressed with PfELC and MLC-B, the MLC-B in the purified myosin is shorter than full-length. Mass spectrometry analysis showed that the cleavage occurred after residue 420. Based on AlphaFold predictions, the large N-terminal region of MLC-B is predominantly α-helical but not involved in PfMyoB binding (**Fig. 1b**). We thus designed a truncated MLC-B that begins at amino acid 421, which removes much of the extended helical region but retains the CaM-like region that binds the heavy chain. This construct was used for co-expression, enabling *in vitro* characterization of a full-length PfMyoB motor bound to both of its light chains, in contrast to a previous study that examined the PfMyoB heavy chain in complex only with MLC-B^15^.

Interestingly, co-expression experiments revealed that PfELC binds to and co-purifies with the FLAG-tagged PfMyoB heavy chain only when MLC-B is also co-expressed (**Fig. 1c**). Thus, PfELC binding depends on the presence of MLC-B, mirroring observations with PfMyoA where PfELC binding requires the presence of MTIP^6^. In contrast, MLC-B and MTIP can each bind to their respective heavy chains independently of the presence of PfELC.

**Figure 1.**
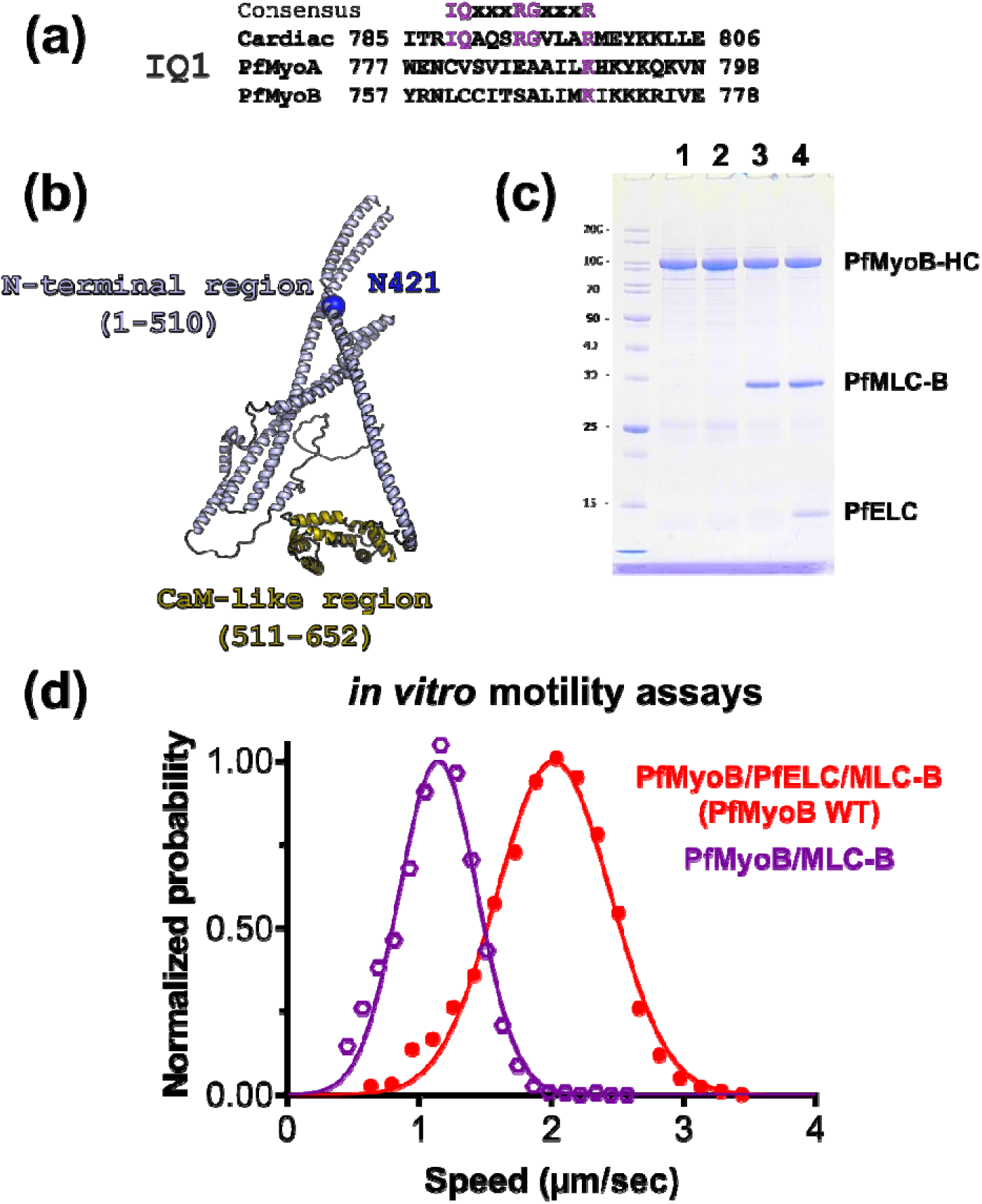
PfELC associates with PfMyoB. **(a)** Sequence alignment of IQ1 from human cardiac myosin (MYH7), PfMyoA and PfMyoB. The consensus sequence is shown and colored in purple. **(b)** AlphaFold model of MLC-B. The N-terminal region (residues 1-510) and the calmodulin (CaM)-like region (residues 511-652) are colored distinctly. **(c)** Coomassie stained 12% SDS-PAGE gel of PfMyoB heavy chain and PUNC co-expression (lane 1). When the PfELC light chain alone is co-expressed with the PfMyoB heavy chain and PUNC (lane 2), the affinity of PfELC for the PfMyoB heavy chain is not enough to co-purify with the PfMyoB heavy chain. The MLC-B light chain co-purifies when co-expressed with PfMyoB heavy chain and PUNC (lane 3). PfELC only binds and co-purifies when co-expressed with the PfMLCB light chain, PUNC and the PfMyoB heavy chain (lane 4). **(d)** Speed distributions from a representative in vitro motility assay of PfMyoB expressed with both PfELC and MLC-B light chains (PfMyoB WT, red) and PfMyoB expressed with only the MLC-B light chain (purple). PfMyoB/PfELC/MLC-B, 2.02 ± 0.43 µm/s; PfMyoB/MLC-B, 1.15 ± 0.30 µm/s. Values reported are mean ± SD. 5239 filaments counted (n=6 movies) for PfMyoB/PfELC/MLC-B and 6686 filaments counted (n=6 movies) for PfMyoB/MLC-B.

To assess light chain occupancy, SDS-PAGE densitometry profiles of PfMyoB and PfMyoA containing their respective co-expressed light chains were compared (**Supplementary Fig. 2c**). For both myosins, the regulatory type light chain (MLC-B or MTIP) exhibited greater Coomassie staining than expected based on their molecular mass. Nevertheless, the relative staining intensities of light chains: heavy chain were similar for the two myosins. Because the crystal structure of PfMyoA demonstrated the binding of one PfELC and one MTIP per heavy chain^6,17^, we infer that PfMyoB likewise binds a single PfELC and a single MLC-B per heavy chain.

The *in vitro* motility speed of PfMyoB increases from 1.15 ± 0.30 µm/s with only MLC-B bound to 2.02 ± 0.43 µm/s with both MLC-B and PfELC bound (**Fig. 1d**). This result further supports PfELC as a functional light chain that increases actin displacement by rigidifying and effectively lengthening the lever arm of myosin, as previously demonstrated with PfMyoA^6^ and class II myosins^18^ containing one versus two light chains.

### Structure of PfMyoB in the Rigor-like state

Extensive crystallization trials were performed with both the full-length (FL) and motor domain constructs (PfMyoB-MD, residues 1-D748, truncated at the end of the Converter) of PfMyoB. While the FL construct was recalcitrant to crystallization, we determined the crystal structure of the MD at 2.0 Å resolution in the nucleotide-free (NF) condition (**Supplementary Table 2, Fig. 2a**). As expected under NF condition, the motor is trapped in a Rigor-like conformation with the Converter and the lever arm swung down and both the actin binding cleft and the inner cleft closed^2,19^(**Fig. 2a; 2b; Supplementary Fig. 3**). The structure aligns with the Rigor-like state of PfMyoA with a RMSD of 1.5 Å. The major difference lies in the angle of the Converter which diverges by ∼28° between the two structures (**Fig. 2b**).

As predicted from sequence alignments, PfMyoB and PfMyoA share several sequence adaptations in their allosteric connectors, including (i) the specific serine at the Fulcrum that restrains the mobility of the SH1-helix and (ii) a degenerate “Wedge” connector lacking the key-aromatic residues (**Fig. 2c, 2d, Supplementary Fig. 1d**). The N-terminal extension (Nterm-extension) is unambiguously resolved in the electron density map (**Fig. 2e**). While the sequence of the N-term extensions of PfMyoA and PfMyoB are highly divergent and do not align properly, both of these extensions contain phosphorylatable residues: S19 for PfMyoA and S16 and T17 in PfMyoB (**Fig. 2d**). Databases such as PlasmoDB (https://plasmodb.org/)^20^ indicate that S16 and T17 are phosphorylated in several stages of the parasite, including the erythrocytic stages. Nonetheless, a kinase that can phosphorylate these sites is not present in *Sf*9 cells, and as isolated neither S16 nor T17 of the heavy chain is phosphorylated in purified PfMyoB. This is in contrast to expressed PfMyoA, which was purified from *Sf*9 cells with S19 phosphorylated^9^. Consistent with the mass spectrometry data, no electron density for phosphorylation at these residues was observed in the PfMyoB structure (**Fig. 2e**).

**Figure 2.**
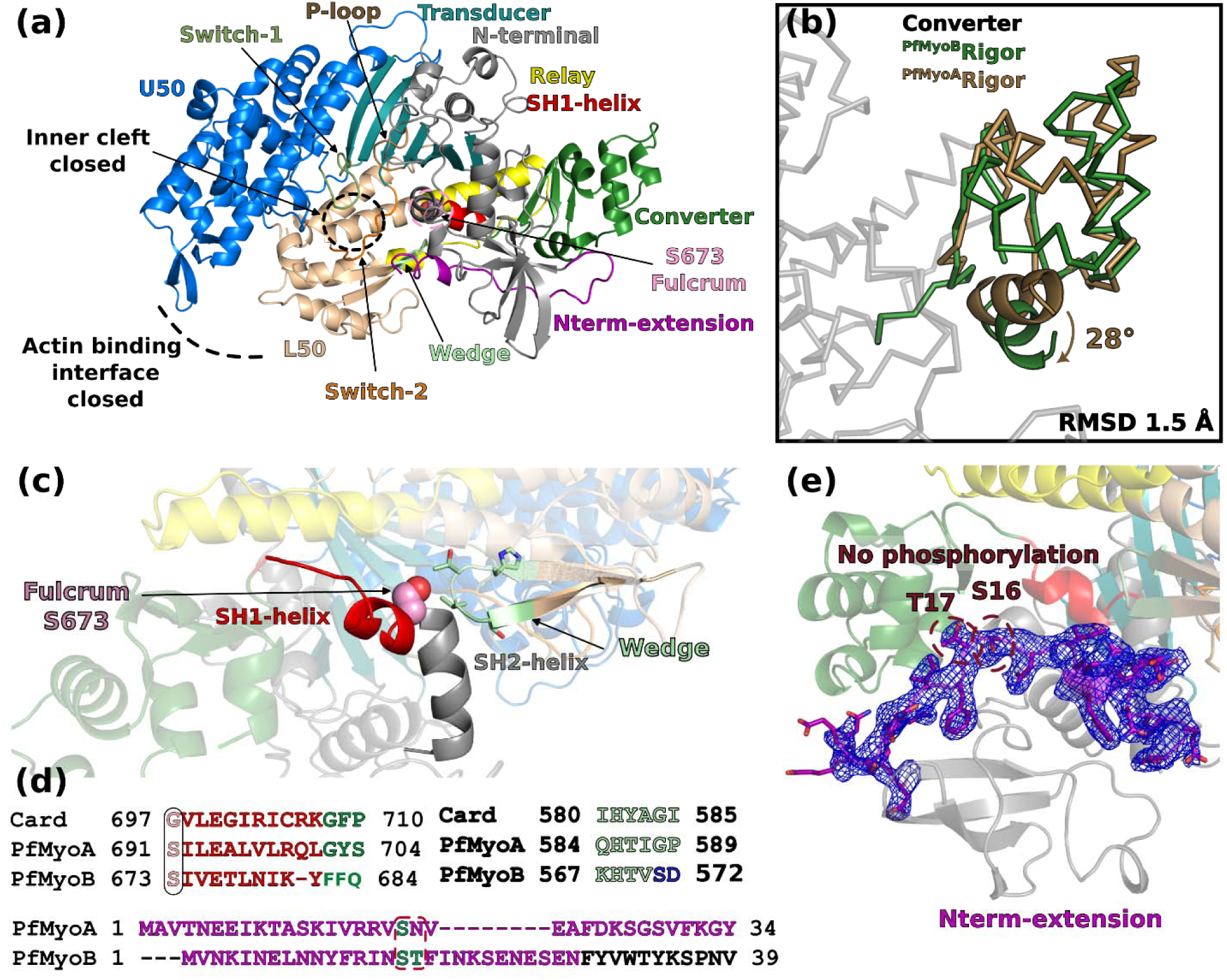
Structure of Plasmodium falciparum Myosin B (PfMyoB). **(a)** Crystal structure of PfMyoB in the Rigor-like state solved at 2.0 Å resolution. Domains and allosteric connectors are colored distinctly. **(b)** PfMyoA Rigor-like structure (6I7D, chain D^9^) has been superimposed on PfMyoB Rigor-like structure (on the entire motor domain). The two structures are highly superimposable with a RMSD of 1.5 Å, the only notable difference being the angle of the Converter that differs ∼28° between the two motors. **(c)** PfMyoB displays atypical sequence differences in the connectors that were previously described in PfMyoA^9^. The most notable is serine 673 instead of the highly conserved glycine at the “Fulcrum”. Other differences are located at the Wedge, where a conserved aromatic is missing. **(d)** Sequence alignment of human cardiac myosin (MYH7, Card), PfMyoA, and PfMyoB for the SH1-helix, the Wedge and the N-terminal extension (Nterm-extension). Residues significantly differing between PfMyoA and PfMyoB in the Wedge are colored in deep blue. Phosphorylatable residues of the Nterm-extension are colored in cyan and contoured in red. **(e)** The N-terminal extension is rebuilt without ambiguity in the electron density map (2Fo-Fc, 1σ). No phosphorylation is seen on S16 or T17.

### The PfMyoB double phospho-mimetic mutant does not enhance *in vitro* motility speed

Phosphorylation of PfMyoA at S19 (SEP19) doubles the speed at which it moves actin in an *in vitro* motility assay relative to a phospho-null S19A mutant^9^. A phosphomimetic S19E mutant also increases motility, although to a lesser extent (∼40%) than bona fide phosphorylation^9^. We therefore hypothesized that if phosphorylation/negative charge similarly regulates PfMyoB activity, introduction of the phosphomimetic mutations S16D /T17E would increase the speed at which PfMyoB moves actin. Although _PfMyoA_K764, which interacts with _PfMyoA_ SEP19, is not conserved in PfMyoB, it aligns with _PfMyoB_H744 and lies in close proximity to _PfMyoB_K740. Due to the orientation of the Converter, these residues could potentially establish interactions with phosphorylated _PfMyoB_S16 and _PfMyoB_T17 in the N-terminal extension of PfMyoB (**Fig. 3a**). However, the double phosphomimetic PfMyoB(S16D/T17E) did not exhibit increased motility relative to unphosphorylated PfMyoB (**Fig. 3b**). These results suggest that, unlike PfMyoA, the introduction of negative charges within the N-terminal extension does not tune the motor’s ability to move actin at different speeds.

### ADP release is not rate limiting for PfMyoB

High duty ratio myosins typically have a limiting ADP-release rate and spend a large fraction of their cycle strongly bound to actin. Conversely, low duty ratio motors are limited by phosphate release rate and spend most of their cycle in detached or weakly actin-bound states (reviewed by^21^). In PfMyoA, S19 phosphorylation modulates the duty-ratio of the motor: although the ADP release rate of the PfMyoA-S19A mutant is not rate-limiting, it is significantly reduced compared to phosphorylated PfMyoA, resulting in slower motility^9^. Interestingly, the ADP-release rate of PfMyoB is more than 4-fold faster compared to its maximum cycling rate in steady-state ATPase assays (474 ± 39 s^-1^ vs. 108 ± 3 s^-1^) (**Fig. 3c**, **Table 1**), indicating that ADP release is not the limiting step in the PfMyoB ATPase cycle (**Fig. 3c**). Additionally, ADP release is unaffected in the phosphomimetic PfMyoB(S16D/T17E) (**Table 1**).

The PfMyoB slow ATPase cycling rate and fast ADP release rate would predict that PfMyoB is a low duty ratio, low force generating myosin. To test this prediction, we measured the relative ensemble force output of PfMyoA and PfMyoB using *in vitro* motility mixture experiments. This approach assesses changes in actin filament velocity as the proportion of two myosins with different speeds is varied^22,23^. The downward curvature observed when phosphorylated PfMyoA is paired with either PfMyoA(S19A) or PfMyoB (**Fig. 3d**) indicates that both PfMyoB and the phosphonull PfMyoA(S19A) mutant have higher ensemble force generating capacity than phosphorylated PfMyoA. Consistent with this result, we previously showed that PfMyoA(S19A) generates higher ensemble force than phosphorylated PfMyoA, using a motility assay in which utrophin acts as a load against which the motor works^9^. The double phosphomimetic mutant PfMyoB(S16D/T17E) shows the same relative ensemble force as unphosphorylated PfMyoB, indicating that negative charges do not modulate its force output. Together with the motility data, these results -show that heavy chain phosphorylation has no measurable effect on PfMyoB motor properties. Contrary to expectations, these assays show that PfMyoB is in fact a high force generating motor.

**Figure 3.**
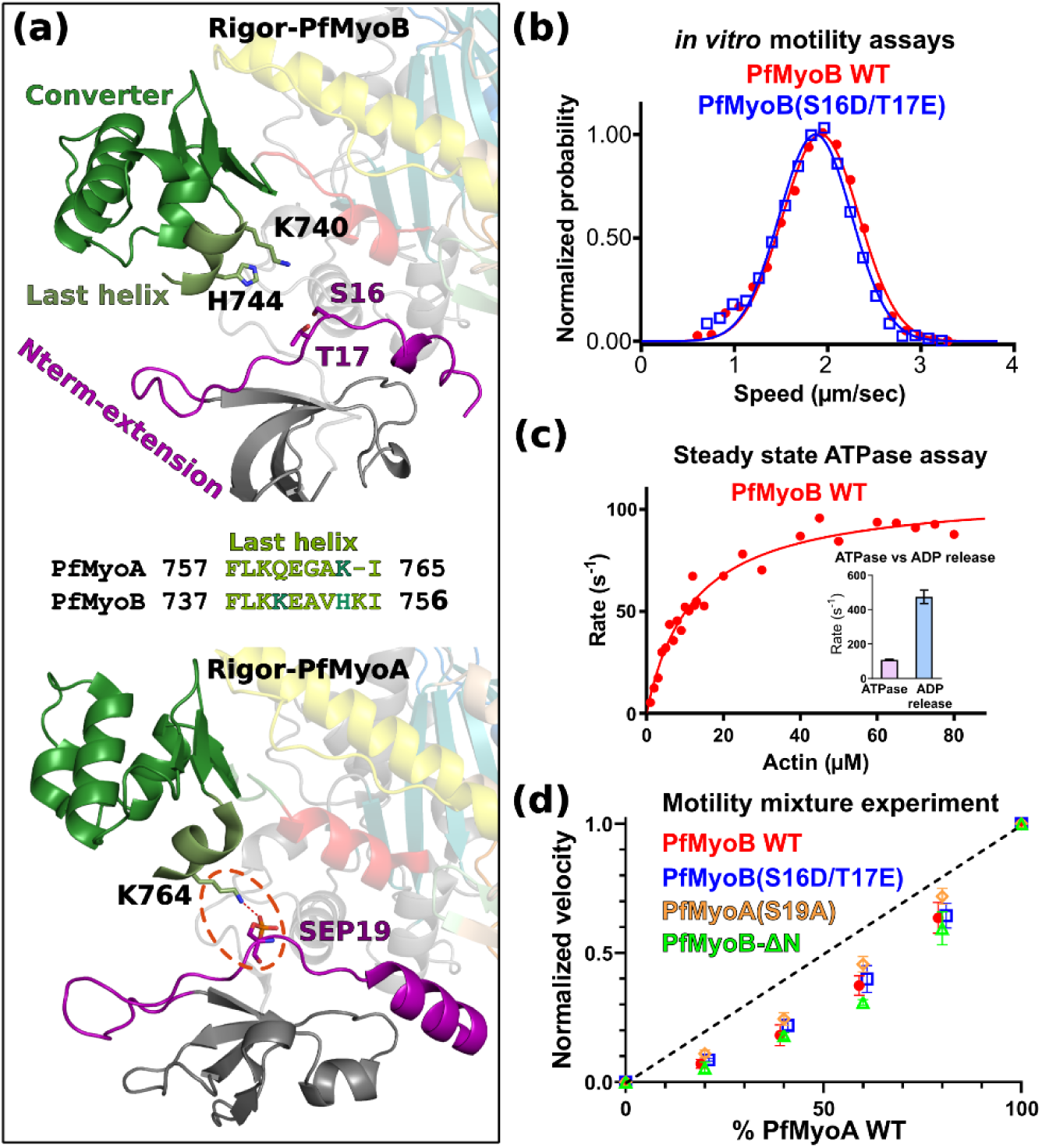
PfMyoB functional properties are not tuned by phosphomimetic mutations. **(a)** shows the potential interactions between the last helix of the Converter and the N-terminal extension (Nterm-extension) in PfMyoB (on the top) and PfMyoA (on the bottom, PDB code 6I7D, chain D^9^) in the Rigor-like state. In PfMyoB, there is no interaction between the Nterm-extension and the last helix of the Converter. In PfMyoA, an electrostatic bond between phosphoserine 19 (SEP19) and K764 stabilizes the Rigor state (contoured with red dashes). In the center, sequence alignment of the last helix of the Converter between PfMyoA and PfMyoB, residues that might interact with phosphorylated S16 and T17 in PfMyoB are colored in dark green. **(b)** Speed distributions from a representative in vitro motility assay of the PfMyoB WT (red) and the phosphomimetic PfMyoB(S16D/T17E) double mutant (blue). PfMyoB WT, 2.02 ± 0.43 µm/s (also shown in Figure 1d); PfMyoB(S16D/T17E), 1.96 ± 0.41 µm/s. Values reported are mean ± SD. 2630 filaments counted (n=6 movies) for PfMyoB(S16D/T17E). The speeds of PfMyoB WT and PfMyoB(S16D/T17E) are statistically insignificant using a two-sided z-test (p=0.9196). **(c)** Actin-activated ATPase activity of PfMyoB WT. V_max_ = 108 ± 3 s^-1^ and K_m_ = 11.9 ± 1.1 mM. Data from two protein preparations and three experiments were fitted to the Michaelis-Menten equation. Error, SE of the fit. **(c inset)** Rate of MyoB WT ATPase vs ADP release rate (at 30°C). **(d)** In vitro motility mixture experiment. The total [myosin] was held constant at 120 µg/ml, while the proportion was varied between PfMyoA WT and PfMyoB (red), PfMyoA WT and MyoB(S16D/T17E) (blue), PfMyoA WT and PfMyoA(S19A) (orange) and PfMyoA WT and PfMyoB-ΔN (green). Data from two protein preparations and three experiments are shown. Temperature, 30°C.

Our results reveal major kinetic differences between PfMyoA and PfMyoB. In PfMyoA, dephosphorylation increases ensemble force at the expense of speed, resulting in a lower ADP release-to-ATP cycling ratio (**Table 1**, data from^9^). Specifically, dephosphorylation reduces the ADP-release rate, increasing the fraction of the cycle spent in strongly actin-bound states and enhancing ensemble force production. In contrast, PfMyoB displays a markedly different behavior. Its ADP release rate is approximately 4-fold higher than that of PfMyoA(S19A), yet both motors exhibit similar and higher ensemble force output compared to phosphorylated PfMyoA. These findings demonstrate that PfMyoB is a high ensemble force generating motor that spends higher fraction of its cycle in a strong actin binding state than would be implied from its fast ADP release kinetics.

**Table 1.** ADP release rates from acto-PfMyoB and acto-PfMyoA at various temperatures. PfMyoB WT, PfMyoB(S16D/T17E) and PfMyoB-ΔN data represent at least three experiments and two protein preparations. PfMyoA WT and PfMyoA(S19A) data are from^9^. Values, mean ± SD.

| **construct** | **20°C** | **25°C** | **30°C** |
| --- | --- | --- | --- |
| **PfMyoB WT** | 72.3 ± 12.0 | 188.3 ± 16.6 | 474.0 ± 39.3 |
| **PfMyoB(S16D/T17E)** | 69.6 ± 2.7 | 179.3 ± 4.5 | 462.7 ± 27.5 |
| **PfMyoB-ΔN** | 23.2 ± 2.6 | 44.6 ± 9.1 | 92.2 ± 10.7 |
| **PfMyoA WT\*** | 103.2 ± 20.1 | 209.0 ± 39.3 | 334.3 ± 36.2 |
| **PfMyoA(S19A)\*** | 33.0 ± 5.3 | 65.7 ± 10.3 | 115.8 ± 10.9 |

### The Nterm-extension stabilizes the Rigor state of PfMyoB

A possible explanation for the high force production of PfMyoB is that the motor spends a larger fraction of its cycle in the strong actin binding Rigor state, which follows ADP release and corresponds to the nucleotide-free actin-bound state. This state is typically transient in myosins, as the active site adopts a conformation promoting rapid ATP rebinding with a closed backdoor^2,24,25^. This, in turn, triggers dissociation of the motor from the actin filament and initiates a new motor cycle^1,26,27^.

**Figure 4.**
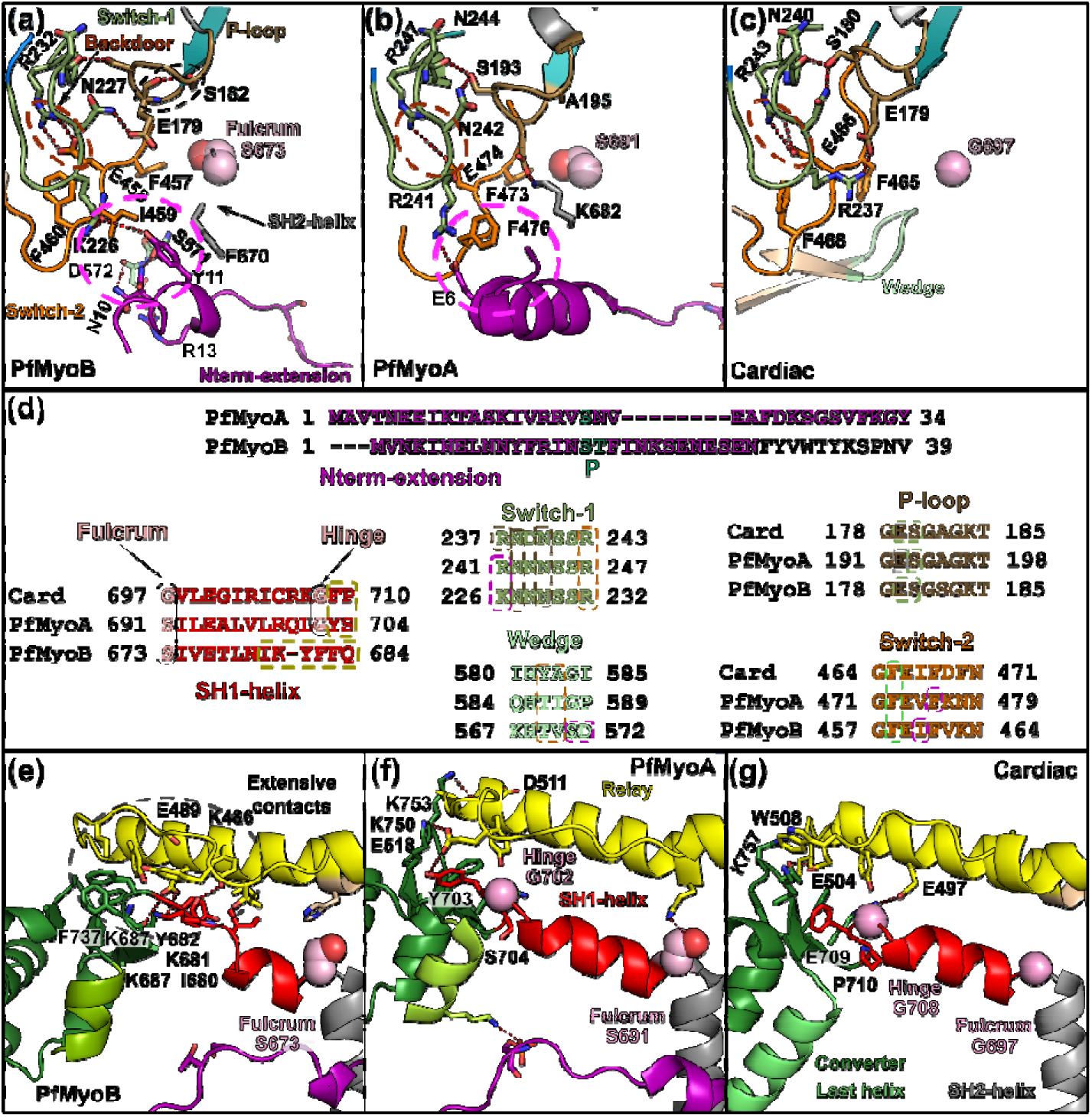
Stabilization of the Rigor state in PfMyoB. **(a-c)** Show the active sites of PfMyoB, PfMyoA (PDB code 6I7D, chain D) and β-cardiac myosin (PDB code 8EFI) respectively, all in the Rigor state. **(d)** Displays a sequence alignment of connectors within the motor domain. Residues making interactions are boxed with dashed lines colored depending on the connector involved in the interaction. **(e-g)** Show the interactions between the SH1-helix/Converter junction and the Relay-helix in PfMyoB, PfMyoA and β-cardiac myosin in the Rigor state.

Interestingly, structural analysis reveals how the Rigor-like conformation is unusually stabilized in PfMyoB. <u>First, allosteric communication is altered (</u>**Fig. 4a-c**). The sequence of the **P-loop** deviates from the highly conserved consensus, featuring a serine (_PfMyoB_S182) instead of the canonical alanine (-GESG**A**GKT-) (**Fig. 4d**). The serine forms a polar interaction with _PfMyoB_E279, maintaining the P-loop in a conformation distinct from PfMyoA and β-cardiac muscle Myo2 heavy chain (βMHC) (**Fig. 4a-c**). Moreover, the **N-term extension** of PfMyoB establishes extensive interactions with multiple allosteric connectors of the motor, including Switch-2, the Wedge and the SH1- and SH2-helices (**Fig. 4a**). This contrasts with PfMyoA, where the N-terminal extension forms only modest interactions with Switch-1 and Switch-2 to maintain the backdoor closed^9^ and with βMHC, where no Nterm-extension interacts with the active site (**Fig. 4b,4c**). These specific structural features are predicted to stabilize the active site in a conformation that slows ATP rebinding.

Another region contributing to stabilization is located at the **base of the Converter** (**Fig. 4e-g**). In most myosins, the Converter base contains a flexible “hinge” with a conserved glycine that enables the Converter swing^28–30^. This glycine is absent in PfMyoB, where this region is enriched in aromatic residues (**Fig. 4e-g**). Specifically, the segment between the SH1 helix and the Converter (residues 679-IKYFF-683) establishes extensive hydrophobic and π–π interactions with the Relay-helix (**Fig. 4f**). This interface is expected to (i) stabilize the orientation of the Converter in the Rigor state and (ii) to slow the reorientation of the Converter due to the absence of glycine.

Collectively, these features likely stabilize the Rigor state of PfMyoB by strengthening interactions involving the Switches and the Wedge, thereby maintaining the Converter in the Rigor orientation. Consequently, nucleotide affinity is reduced, as the nucleotide binding pocket must disrupt multiple stabilizing interactions to adopt a conformation capable of tightly binding nucleotide.

### Weak ATP affinity and slow ATP binding result in more ATPase cycling time spent in a strongly bound to actin state

To further test our hypothesis that PfMyoB has weak nucleotide affinity caused by the unique PfMyoB N-term extension stabilizing interactions, we performed *in vitro* motility speed assays over a range of ATP concentrations. If true, the maximum *in vitro* motility speeds for MyoB would be achieved at higher ATP concentrations than PfMyoA, which has an N-terminal extension that makes fewer stabilizing contacts with the nucleotide binding pocket, and secondly, the maximum *in vitro* motility speed for PfMyoB would be independent of phosphorylation state, as assessed by the phosphomimetic PfMyoB(S16D/T17E) construct. Indeed, PfMyoB-WT and PfMyoB(S16D/T17E) exhibit a ∼4-fold weaker K_M_ for ATP than phosphorylated PfMyoA and PfMyoA(S19A) in the presence of actin (**Fig. 5a**). Furthermore, the rate of ATP binding to PfMyoB or PfMyoB(S16D/T17E) is considerably slower than that of phosphorylated PfMyoA or PfMyoA(S19A) in the presence of actin, indicating that the PfMyoB nucleotide binding pocket cannot adopt a conformation capable of tight ATP binding as readily as PfMyoA (**Fig. 5b**).

We also compared the nucleotide affinity of PfMyoB to PfMyoA in the absence of actin (**Supplementary Table 3, Fig. 5c, 5d**). Fluorescent nucleotide binding experiments show that PfMyoB has a 5-fold weaker ATP affinity than PfMyoA (**Supplementary table 3, Fig. 5c**), resulting from both slower ATP binding and faster ATP release (**Supplementary table 3, Figure 5c**). These two factors can be interpreted in light of the structure: <u>the specific conformation of the PfMyoB active site is less well suited for ATP binding (accelerated ATP release), reducing the probability of inducing a tight ATP binding state (slower ATP binding)</u>. PfMyoB also binds ADP more slowly than phosphorylated PfMyoA in the absence of actin (**Figure 5d**), and its overall ADP affinity is also lower, confirming that the nucleotide-free state of PfMyoB exhibits weaker nucleotide binding affinity (**Supplementary table 3**).

**Figure 5.**
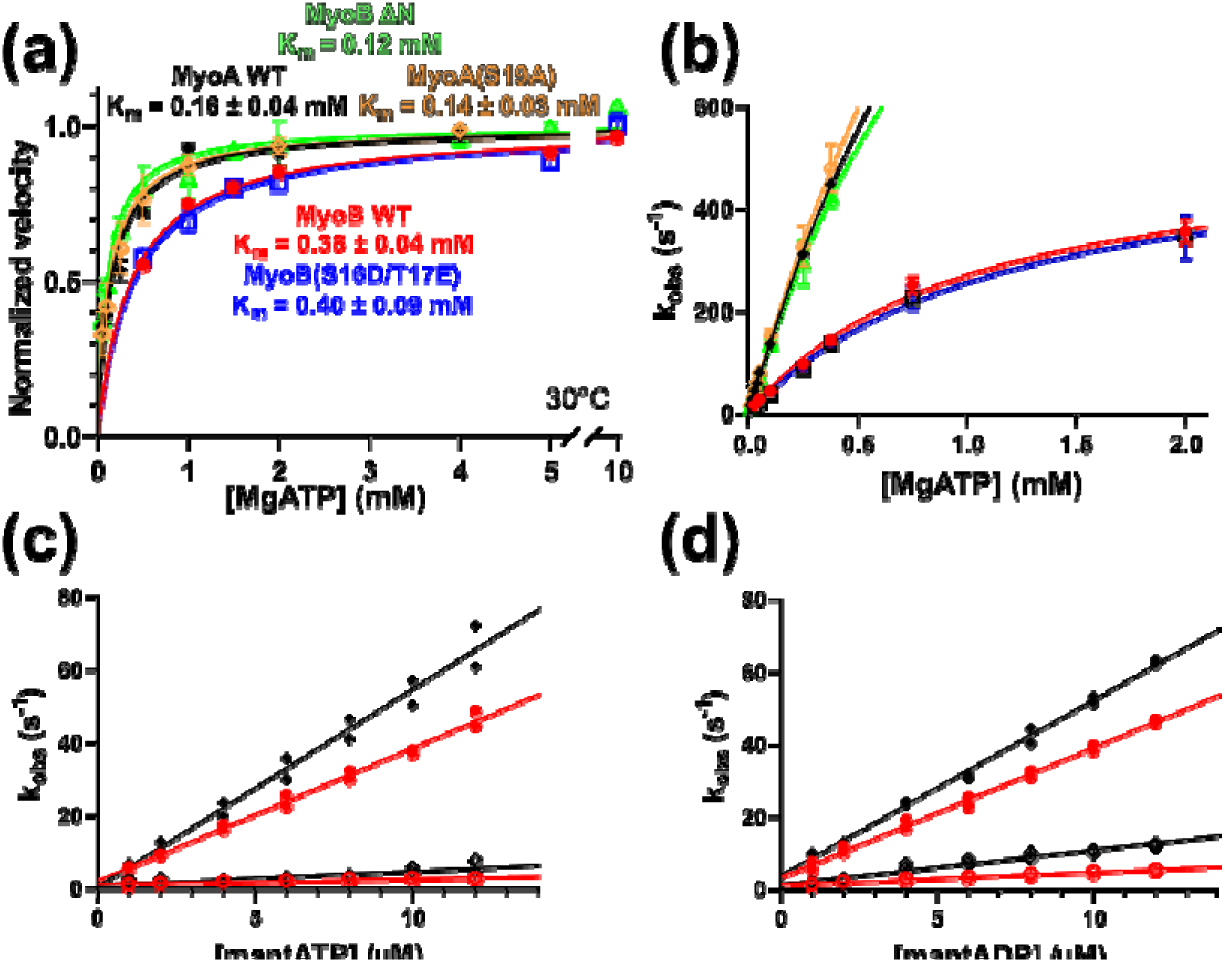
PfMyoB has a weak nucleotide affinity. **(a)** In vitro motility speed with increasing [MgATP] for PfMyoB WT (red), PfMyoB(S16D/T17E) (blue), PfMyoA WT (black), PfMyoA(S19A) (orange) and PfMyoB-ΔN (green). Data from 2 protein preparations and 3 experiments were fitted to the Michaelis-Menten equation. Error, SE of the fit. **(b)** Rate of actomyosin dissociation by MgATP for the same myosin constructs shown in panel (a). Data from at least 2 protein preparations and 3 experiments were fitted to the Michaelis-Menton equation. Temperature, 15°C. Error, SE of the fit. Nucleotide binding rate for PfMyoB WT (red) and PfMyoA WT (black) measured with mant-ATP **(c)** and mant-ADP **(d)**. PfMyoA WT data from^10^. Data from two experiments with independent protein preparations are shown. The association (slope) and dissociation (y-intercept) rate constants and calculated affinities are tabulated in **supplementary table 3.**

Altogether, these results explain the unusual behavior of PfMyoB. **Its slow and weak ATP binding causes the motor to spend a larger fraction of the cycle in the nucleotide-free state, thereby increasing the duty ratio of PfMyoB despite its very fast ADP-release rate.**

### Deletion of the PfMyoB N-term extension enhances ATP binding properties and slows ADP release

To demonstrate that the slow and weak ATP binding properties of PfMyoB are directly mediated by its N-term extension, we expressed a truncated construct beginning at residue 29 that lacks the N-terminal extension (PfMyoB-ΔN). We predicted that removal of the N-terminal extension would result in tighter and faster ATP binding than observed for WT PfMyoB. Consistent with this prediction, PfMyoB-ΔN exhibited a 4-fold lower K_M_ for ATP (tighter affinity) than WT-PfMyoB, yielding values similar to those measured for PfMyoA (**Fig. 5a**). In addition, ATP binding rates in the presence of actin were greatly accelerated in PfMyoB-ΔN relative to WT-PfMyoB and similar to those of the PfMyoA constructs. We further predicted that the deletion of the N-term extension would strengthen nucleotide binding and therefore slow ADP release. Consistent with this hypothesis, the rate of ADP release from actomyosin was reduced approximately 5-fold in PfMyoB-ΔN compared with WT-PfMyoB (474.0 ± 39.3 s^-1^ to 92.2 ± 10.7 s^-1^) (**Table 1**). Interestingly, the ADP release of PfMyoB-ΔN is close to PfMyoA(S19A) (92.2 ± 10.7 s^-1^ vs 115.8 ± 10.9 s^-1^ at 30°C). This finding suggests that removal the N-term extension of PfMyoB restores a higher affinity for the nucleotides. The fact that PfMyoB-ΔN still generates higher ensemble force than phosphorylated PfMyoA (**Fig. 3d**) is fully consistent with our analysis: although PfMyoB-ΔN has higher ATP affinity than the WT and detaches rapidly from F-actin, its slower ADP release rate results in higher ensemble force that is similar to unphosphorylated PfMyoA(S19A). This behavior is also consistent with both myosins sharing a serine instead of a glycine at the Fulcrum, preventing piston-like movement of the SH1 helix, a characteristics that was demonstrated to slow ADP-release in PfMyoA when unphosphorylated^9^.

**Figure 6.**
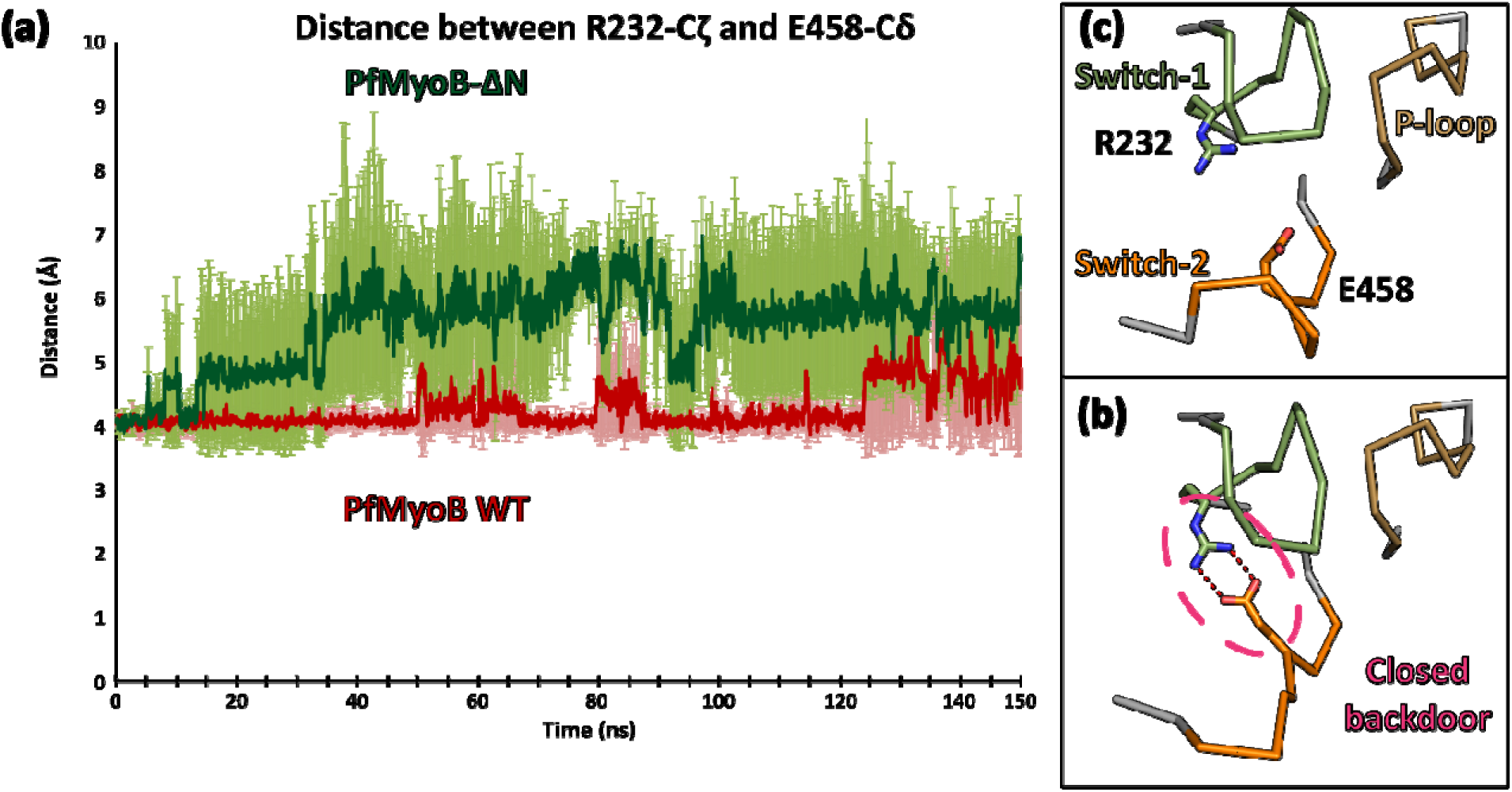
The N-terminal extension of PfMyoB restrains the conformational exploration of the active site. All-atom molecular dynamics experiments were conducted during 150 ns on two conditions: the wild-type motor domain (PfMyoB WT) and a motor domain in which the N-terminal extension was truncated (PfMyoB-ΔN). **(a)** Shows the distance between R232 (Switch-1) and E458 (Switch-2) which form the so-called “backdoor” that can be “closed” when they establish an electrostatic interaction **(b)** or in a more “open” position **(c)**. Each simulation was performed three times and the standard deviation is shown in **(a)**. The distance between these two residues reflects the ability of the backdoor to adopt open or closed states. Interestingly, the backdoor remains in the closed state for longer in PfMyoB WT than in PfMyoB-ΔN, indicating that the N-terminal extension of PfMyoB restricts the conformational dynamics of the backdoor and more broadly, of the active site.

To directly monitor the effects of the N-terminal extension of PfMyoB on its active site, we performed all-atom molecular dynamics on the motor domain in two conditions: PfMyoB-WT and PfMyoB-ΔN (**Supplementary Fig. 4a, Supplementary Movie 1**). Each simulation was run for 150 ns (see Methods), a timescale sufficient to observe mutation- or truncation-induced changes in myosin motors (e.g. ^31^). Each condition was repeated three times to ensure the reproducibility of the findings (**Supplementary Fig. 4b-c**). Overall, the WT condition is more dynamic than ΔN (**Fig. 6a**) and it is mainly due to the first residues of the N-terminal extension that explore different conformations (**Supplementary Fig. 4b-c**). No obvious differences in the dynamics of the P-loop, Switch-1 and Switch-2 were observed between the PfMyoB-WT and -ΔN based on the RMSF curves (**Supplementary Fig. 4b-c**). However, a detailed inspection of the trajectories reveals that the N-terminal extension of PfMyoB restricts the conformational dynamics of both Switches-1 and -2, as these regions can explore distinct conformations when the N-terminal extension is deleted (**Supplementary Movie 2**). More specifically, closure of the “backdoor” can be monitored by measuring the distance between _Switch-1_R232 and _Switch-2_E458 (**Fig. 6a**). Interestingly, the N-terminal extension limits the flexibility of Switches-1 and -2, maintaining the backdoor in a closed conformation (**Fig. 6b**) for most of the PfMyoB WT simulations (**Fig. 6a**). In PfMyoB-ΔN, the backdoor shows a greater propensity to lose this electrostatic interaction (**Fig. 6c**) and to explore alternative conformations (**Fig. 6a, Supplementary Movie 2**). These findings demonstrate that the N-terminal extension of PfMyoB restricts backdoor dynamics, favoring a closed conformation.

Altogether these results support the hypothesis that the N-terminal extension from PfMyoB restrains the conformational changes of the active site connectors, which reduces nucleotide affinity. Consistent with the *in vitro* assays, structural and dynamics experiments reveal that the extensive interactions of the N-terminal extension have two major effects: (i) they accelerate the ADP release from the motor by stabilizing a unique low nucleotide affinity conformation and (ii) delay ATP binding in the Rigor state, resulting in a motor that spends a larger fraction of its mechanochemical cycle in a nucleotide-free state.

**Figure 7.**
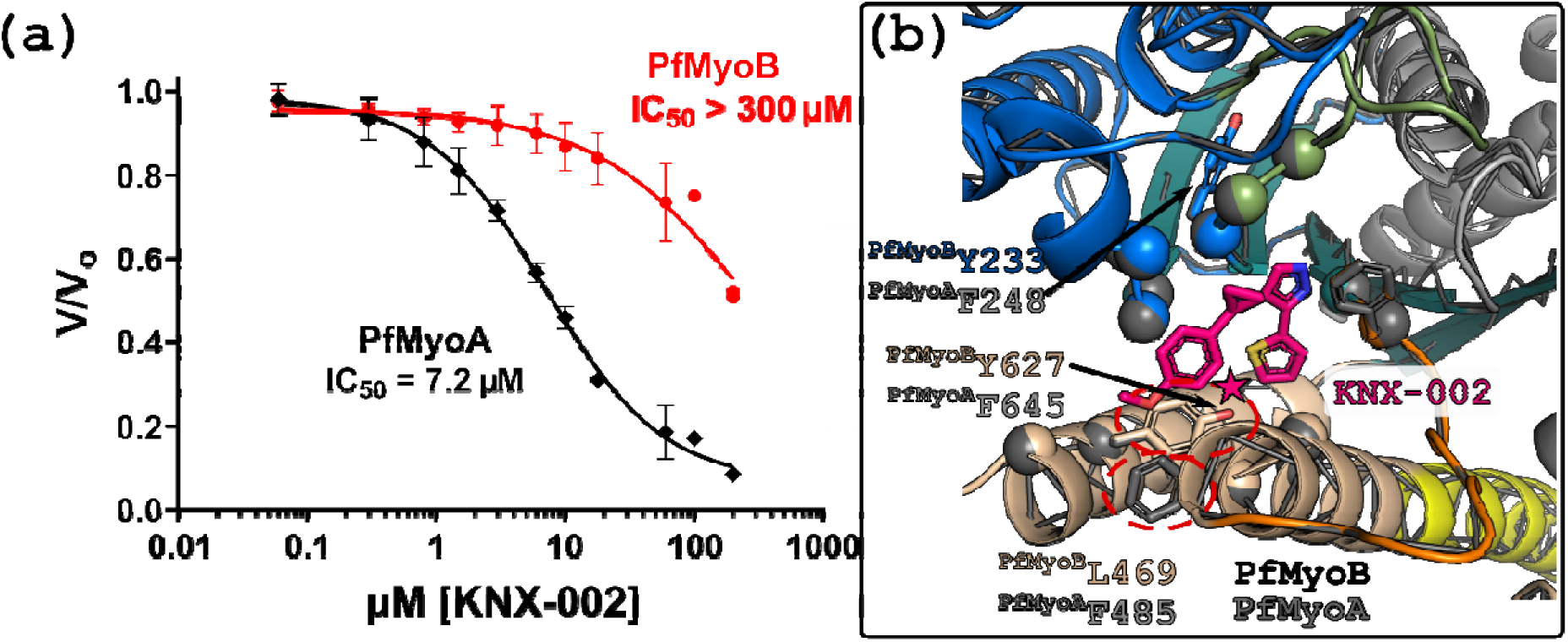
The activity of PfMyoB is weakly inhibited by KNX-002. **(a)** Dose-response curve measuring the ATPase activity of PfMyoA and PfMyoB in presence of actin with increasing concentrations of KNX-002 (50 µM actin, 30°C). The ATPase activity of PfMyoB is weakly inhibited by KNX-002 relative to the previous measurement of KNX-002 inhibition of PfMyoA^10^. **(b)** Homology model of PfMyoB (PfMyoB) generated with PfMyoA bound to KNX-002 as template (PfMyoA, PDB code 8CDQ). The KNX-002 binding pocket in PfMyoB shows three differences compared to PfMyoA: ^PfMyoA^F248 is replaced by ^PfMyoB^Y233; ^PfMyoA^F485 is replaced by ^PfMyoB^L469 resulting in a loss of a π-stacking interaction and ^PfMyoA^F645 is replaced by ^PfMyoB^Y627, introducing steric hindrance with the compound (pink star). The differences at positions 485 and 645 can explain why KNX-002 is less efficient on PfMyoB compared to PfMyoA.

### KNX-002 exerts a weak inhibition on PfMyoB activity

KNX-002 is a small molecule recently identified as an inhibitor of PfMyoA ATPase activity, as well as parasite pathogenesis^10,32^. Since PfMyoA and PfMyoB are both class XIV myosins, we tested whether KNX-002 could also inhibit PfMyoB ATPase activity despite sequence divergence between the two motors. Interestingly, KNX-002 weakly inhibits PfMyoB compared to PfMyoA (IC_50_ >300 vs 7.2 µM, **Fig. 7a**).

To understand this difference, we generated a homology model of PfMyoB bound to KNX-002 using the PfMyoA/KNX-002 structure (PDB code 8CDQ^10^) as a template (**Fig. 7b**) Most residues involved in KNX-002 binding are conserved between PfMyoA and PfMyoB, with three notable exceptions: ^PfMyoA^F248 is replaced by ^PfMyoB^Y233; ^PfMyoA^F485 is replaced by ^PfMyoB^L469 and ^PfMyoA^F645 is replaced by ^PfMyoB^Y627. While ^PfMyoA^F248/^PfMyoB^Y233 substitution is predicted to have little effect on KNX-002 binding, ^PfMyoA^F485/^PfMyoB^L469 and ^PfMyoA^F645/^PfMyoB^Y627 substitutions are likely to have more severe consequences. Specifically, ^PfMyoA^F485/^PfMyoB^L469 disrupts a key π-stacking interaction with the methoxyphenyl group of KNX-002, and ^PfMyoA^F645/^PfMyoB^Y627 introduces a steric hindrance with the thiophene group (**Fig. 7b**). These two significant sequence differences in the KNX-002 binding pocket likely explain the reduced efficacy of the compound against PfMyoB by weakening its binding affinity.

## Discussion

Despite both PfMyoB and the previously characterized PfMyoA^9,17,33^ being class XIV *Plasmodium* myosin motors, they differ markedly in both sequence and structure. By integrating structural and kinetic analyses, we show that these motors employ distinct mechanochemical strategies (**Figure 8a-b**). PfMyoA is a tunable motor whose duty-ratio is modulated by phosphorylation of Ser19 in its N-terminal extension (**Fig. 8a**). In contrast, introducing negative charge in the N-terminal extension of PfMyoB (S16D/T17E) has no effect on its kinetic or motile properties, and the motor appears to be specialized for slower speed and high ensemble force generation (**Fig. 8b**). Moreover, the N-terminal extension of PfMyoB tunes motor properties through a markedly different allosteric mechanism compared to PfMyoA. PfMyoA has a tunable mechanism of force production in which the N-terminal extension accelerates the ADP-release rate of the motor when phosphorylated. In contrast, PfMyoB stabilizes the nucleotide-free state through three principal adaptations. First, the active site is maintained in a conformation that is incompatible with fast ATP binding due to stabilizing interactions between the N-terminal extension and Switch-1, Switch-2 and the Wedge. Second, substitution of a canonical alanine with a serine alters the conformation of the P-loop. Third, sequence divergence at the base of the Converter stabilizes the rigor conformation (**Fig. 8b**). Remarkably, deletion of the N-terminal extension of PfMyoB is sufficient to change the affinity and kinetics of nucleotide binding, resulting in faster binding of ATP to acto-PfMyoB, tighter ATP binding and slowed ADP release. These mechanistic differences are consistent with the distinct cellular functions of the two motors. PfMyoA is part of the glideosome and contributes both to parasite motility that requires speed, and to red blood cell invasion that requires force production against load. In contrast, PfMyoB localizes to the apical region and appears to function specifically during the initiation of invasion, a process that likely depends on sustained force generation (**Fig. 8a-b**).

Interestingly, both PfMyoB and PfMyoA bind the same essential light chain PfELC^6,7^, despite binding distinct light chains at their second IQ motifs (MLC-B and MTIP, respectively). Previous structural studies of full-length PfMyoA in complex with PfELC and MTIP demonstrated that PfELC binds to its target IQ motif (IQ1) in an unconventional manner relative to canonical CaM-like light chains, because of the degenerate sequence of both PfELC and IQ1^17^. Our finding that IQ1 from PfMyoB (_PfMyoB_IQ1) also associates with PfELC despite strong sequence divergence from _PfMyoA_IQ1 is therefore particularly intriguing. In PfMyoA, W777 inserts into the C-lobe of PfELC^17^, whereas it is replaced by Y757 in PfMyoB. These observations suggest that PfELC may engage in distinct interaction modes with PfMyoA and PfMyoB, potentially giving rise to lever arms with different structural and mechanical properties.

The mechanisms by which PfMyoA and PfMyoB tune their motor properties can be compared to those described for class I myosins. In myosin-1b (Myo1b), the duty ratio is strongly regulated by mechanical load, whereas myosin-1c (Myo1c) is largely force-insensitive^34–36^. As a result, Myo1b functions as a force-sensitive membrane-actin anchor, while Myo1c acts as a more dynamic actin-membrane tether. Like PfMyoA, class-I myosins use their N-terminal extensions to modulate allosteric communication between the active site and the Converter (**Fig. 8c**). In Myo1b and Myo1c, distinct interactions involving the N-terminal extension give rise to markedly different mechanochemical behaviors^35,36^. In Myo1b, the N-terminal extension forms part of the Converter/motor-domain interface, delaying lever-arm rotation and ADP release under force^34,35^ (**Fig. 8c**). Conversely, in Myo1c the N-terminal extension contributes to the Converter/motor-domain interface specifically in the rigor state, thereby slowing ATP binding rather than ADP release^35,36^. Thus, Myo1b and Myo1c differ in how they couple active-site rearrangements to lever-arm motion. As with Myo1b, direct interactions mediated by the phosphorylatable N-terminal extension of PfMyoA, the Converter and the active site^9^ likely modulate communication between the active site and the lever arm, tuning ADP release and motor properties. The effects of the N-terminal extension differ in PfMyoB and in class I myosins. In PfMyoB, there is likely no effect of the N-terminal extension on the Converter, instead, the main effects are observed on the active site, where it modulates nucleotide affinity.

Allosteric tuning mechanisms have also been described in dimeric processive motors such as myosin-V (MyoV) and myosin-VI (MyoVI). Their processivity relies on gating, whereby mechanical strain slows the leading head, allowing the trailing head to detach from actin and step forward^37–40^ (**Fig. 8d**). Similar to PfMyoB, MyoVI slows ATP rebinding in the rigor state through direct interactions between a unique insertion, called Insert-1, and active site elements^41^. This mechanism selectively reduces the ATP binding rate without significantly affecting ADP release, thereby preventing premature detachment from actin under strain^41^. In contrast, strain in MyoV primarily slows the ADP-release rate^37–40^. Together, these examples illustrate how relatively conserved myosin cores can be functionally diversified through ancillary structural elements and altered allosteric coupling pathways. In many cases, integrative structural biology has been instrumental in uncovering the molecular basis of these adaptations and linking them to specific cellular functions. Delaying either ADP release or ATP binding represents two alternative strategies for increasing myosin duty-ratio. Although the response of PfMyoA and PfMyoB to mechanical strain remains unknown, their markedly different mechanochemical mechanisms, cellular localizations and biological functions strongly suggest distinct force-dependent behaviors. Our study provides a framework for future *in vivo* investigations aimed at understanding the precise cellular role of PfMyoB, a slow motor specialized for reduced ATP-binding kinetics and sustained force generation.

Mechanochemical tuning appears to be particularly important in Apicomplexan parasites, which rely on a limited repertoire of myosins and light chains to fulfill a wide range of cellular functions. Although MyoA activity is regulated by phosphorylation in both *Plasmodium falciparum* and *Toxoplasma gondii*, the two motors have substantially diverged in sequence (∼63% identity). TgMyoA contains three phosphorylatable residues within its N-terminal extension that exert incremental effects on motility when replaced by phospho-mimetic substitutions^42^. Interestingly, *Toxoplasma gondii* does not express MyoB but instead expresses another class-XIV myosin, TgMyoH, which promotes conoid protrusion during the initiation of motility^43^. TgMyoH associates with conoid microtubules through its tail domain, and conditional depletion of this motor severely impairs parasite motility, invasion and egress^43^. These differences likely reflect the distinct organization of the apical complex in *Plasmodium* and *Toxoplasma*. *T. gondii* possesses a prominent conoid composed of helically arranged microtubules that protrudes during invasion and egress^44^. In contrast, the conoid of *Plasmodium* parasites was long thought to be lost but is now recognized as a small remnant structure^45^. These architectural differences may underlie the requirement for distinct myosin repertoires and explain the variable phenotypic consequences associated with their loss. More broadly, the remarkable structural and mechanochemical diversity displayed by the limited set of Apicomplexan myosins highlights how these parasites expand functional output by exploiting flexible interaction modes and tunable force-production mechanisms. Such evolutionary plasticity likely facilitates adaptation to the diverse cellular environments, host tissues and host species encountered throughout their complex life cycles.

With the discovery of the MyoA inhibitor KNX-002, Apicomplexan myosins have emerged as promising antimalarial targets^10,11^. The subsequent characterization of the more potent derivative KNX-115 further demonstrates the feasibility of optimizing the KNX scaffold^11^. Here, we show that despite the strong divergence between PfMyoA and PfMyoB, KNX-002 still retains measurable inhibitory activity against PfMyoB. Although PfMyoB is not essential for parasite survival or pathogenesis, the simultaneous targeting of PfMyoA and PfMyoB may represent an attractive strategy for disrupting multiple motors involved in host cell invasion. More broadly, the remarkable mechanochemical specialization of Apicomplexan myosins, combined with their central roles in parasite motility and invasion, highlights these motors as attractive targets for the development of a new generation of antimalarial compounds. The structural and mechanistic diversity uncovered within this motor family may also provide opportunities for the design of compounds with distinct specificities and modes of action.

**Figure 8.**
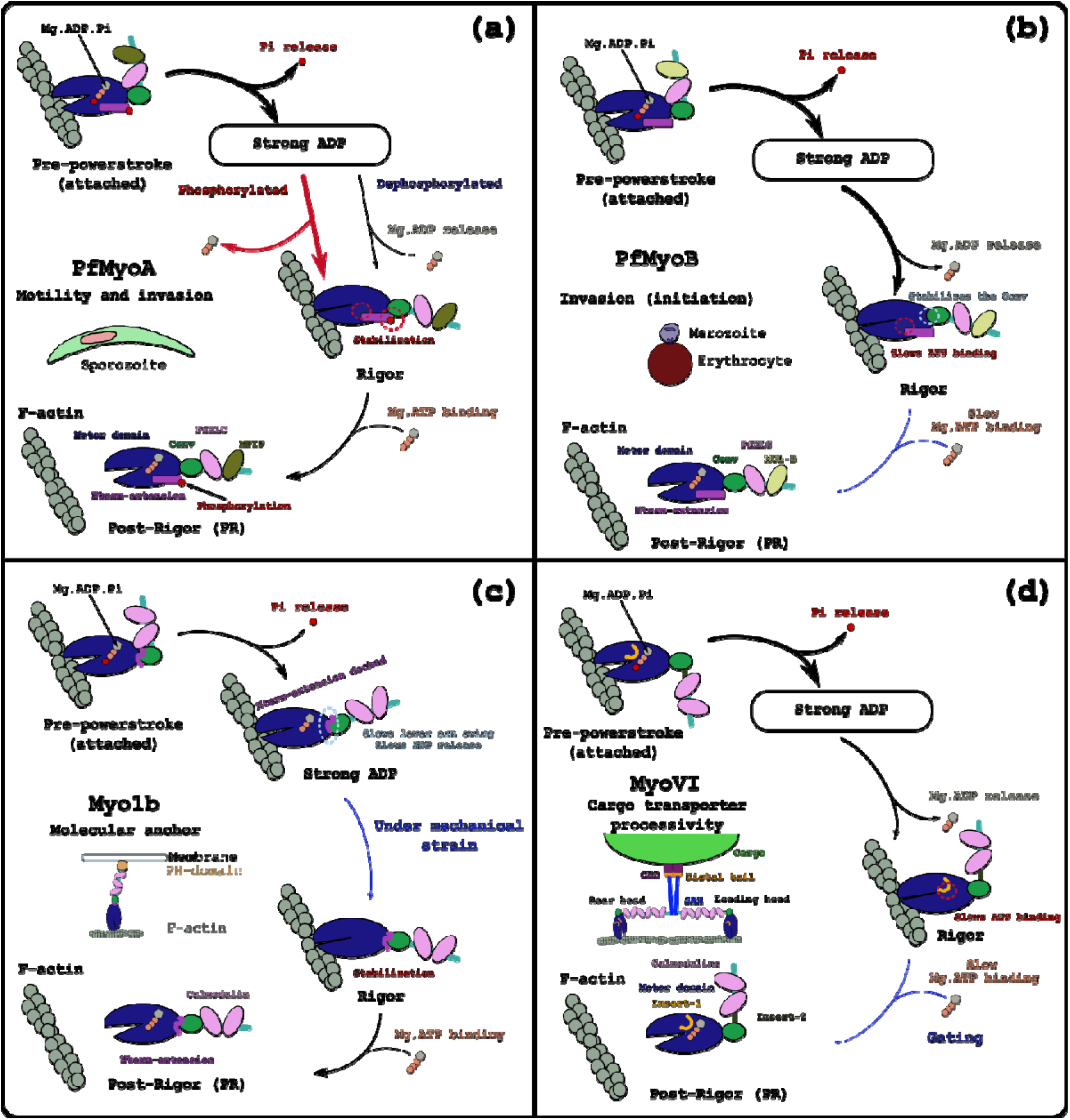
allosteric tuning of myosins. **(a)** PfMyoA has a tunable mechanism of force production. The phosphorylation of the N-terminal extension (Nterm-extension) modulates the Pi release rate. This motor is involved in parasite motility and invasion. **(b)** PfMyoB is a slow motor tuned for force production. The Rigor state is stabilized by (i) specific interactions between the base of the Converter and the motor domain (blue), (ii) the Nterm-extension slowing ATP binding (red). This motor is tuned for ensemble force production. **(c)** Myo1b is a molecular anchor connecting the membrane to cortical actin. Under mechanical strain, interactions between the Nterm-extension and the Converter in the strong ADP state slows both ADP release and lever arm swing, therefore increasing the duty-ratio. PH-domain: pleckstrin homology domain. **(d)** The cargo transporter MyoVI is tuned for processivity. Under mechanical strain, detachment of the leading head is delayed. A specific region called insert-1 interacts with the active site and slows ATP binding (in red). MyoVI is also characterized by insert-2 at the end of the Converter this insert enables the myosin to move in the opposite direction along the actin filament compared with other myosins^1,46^.

## Methods

### Key resources

Key resources used in this study are tabulated in Supplementary Table **X**.

### SDS-PAGE gel quantification

Densitometry analysis was performed on all images using ImageJ. The gel images shown for quantification are uncropped and unmodified.

### Actin-activated ATPase assays

The actin-activated ATPase activity of PfMyoB (20 nM) was assessed at 30°C in a buffer containing 10 mM imidazole (pH 7.5), 5 mM KCl, 1 mM MgCl_2_, 1 mM EGTA, 1 mM DTT, and 1 mM NaN_3_. 2 mM MgATP was and ATPase activity was measured using a linked assay that couples the regeneration of hydrolyzed ATP to the oxidation of NADH. ATPase assays with KNX-002 incorporated 1% DMSO in the buffer. The change in optical density at 340 nm was recorded over time on a Lambda 25 UV/VIS spectrophotometer (Perkin Elmer), with data fitting performed according to the Michaelis-Menten equation (Graph Pad Prism v10.6.1).

### Protein expression and purification

PfMyoB heavy-chain constructs (WT, S16D/T17E and ΔN) were co-expressed with a *Plasmodium spp.* UCS (UNC-45/CRO1/She4p) family myosin chaperone^6^ and the light chains PfMLC-B and PfELC. The full-length WT heavy-chain construct comprises the complete *P. falciparum* myosin B sequence (Plasmo DB PF3D7_0503600 accession number XM__001351558.1), followed by a 13–amino-acid spacer (NVSPATVQPAFGS), an 88–amino-acid fragment of the *E. coli* biotin carboxyl carrier protein^47,48^, and a C-terminal FLAG tag. The PfMyoB(S16D/T17E) and PfMyoB-ΔN (Δ residues 1-27) variants were generated by site-directed mutagenesis using the WT heavy-chain gene as template. For PfMyoA crystallization, a MD construct (residues 1-D748) followed by a 2-amino-acid spacer (GS) and a C-terminal FLAG tag was expressed. PCR products were cloned into the baculovirus transfer vector pAcSG2 (554769, BD Biosciences) to generate recombinant virus. The *Plasmodium* UCS myosin chaperone used for co-expression is a chimeric clone combining the *P. falciparum* (Plasmo DB PF3D7_1420200/GenBank^TM^ accession number XM_001348333.1) coding sequence with regions from *P. knowlesi* (Plasmo DB PKNH_1337800/GenBank^TM^ accession number XM_002260772) at loci lacking *Plasmodium* interspecies consensus. A C-terminal Myc tag was appended and the resulting product cloned into pAcSG2 for recombinant virus production. PfELC (Plasmo DB PF3D7_1017500/GenBank^TM^ accession number XM_001347419.1) and full-length PfMLC-B (PlasmoDB PF3D7_1118700/GenBank^TM^ accession number XM_001347828.2) were amplified from gBlocks® gene fragments (Twist Bioscience) and subcloned into the pFastBac vector (10359016, Thermo Fisher Scientific) for recombinant baculovirus generation. A truncated PfMLC-B (amino acid residues 421–652) was produced by site-directed mutagenesis from the full-length PfMLC-B template.

Sf9 cells (71104, Novagen) were infected with recombinant baculovirus and cultured for 72 h in medium supplemented with 0.2 mg·mL−1 biotin. Cells were harvested and lysed by sonication in lysis buffer composed of 10 mM imidazole, pH 7.4, 0.2 M NaCl, 1 mM EGTA, 5 mM MgCl_2_, 7% (w/v) sucrose, 2 mM DTT, 0.5 mM 4-(2-aminoethyl) benzene-sulfonyl fluoride, 5 μg·mL^−1^ leupeptin, and 2 mM MgATP. An additional 2 mM MgATP was added prior to clarification by centrifugation at 200,000 × g for 40 min. The clarified supernatant was applied to an anti-FLAG M2 affinity gel column (A2220, Sigma-Aldrich). The column was washed with a buffer containing 10 mM imidazole, pH 7.4, 0.2 M NaCl, and 1 mM EGTA, and the bound myosin was eluted with the same buffer supplemented with 0.1 mg·mL^−1^ FLAG peptide (A6002, APExBIO). Myosin-containing fractions were pooled, concentrated using an Amicon centrifugal filter device (901024, Millipore), dialyzed overnight against 10 mM imidazole, pH 7.4, 0.2 M NaCl, 1 mM EGTA, 55% (v/v) glycerol, 1 mM DTT, and 1 μg·mL^−1^ leupeptin, and stored at −20 °C.

Skeletal muscle actin was isolated from chicken pectoralis major obtained from frozen tissue (Trader Joe’s frozen chicken breasts). Muscle tissue was minced with a meat grinder and suspended in extraction buffer (150 mM potassium phosphate pH 6.7, 300 mM KCl, 2 mM EDTA, 1 mM DTT and 3 mM NaN_3_). The homogenate was transferred to a beaker covered with cheesecloth and rinsed with 10 mM sodium bicarbonate containing 0.1 mM CaCl_2_. The tissue was sequentially washed with deionized water and acetone, then air-dried overnight to produce acetone powder. Actin was purified from the acetone powder following established procedures^49^.

Acetone powder was resuspended in G-buffer (5 mM Tris, pH 8.2 at 4 °C; 0.2 mM CaCl_2_; 0.2 mM Na_2_ATP; 0.5 mM DTT; 1 μg·mL^−1^ leupeptin) and G-actin was extracted for 1 h at 4 °C. Insoluble material was removed by centrifugation at 35,000 × g for 20 min. The clarified G-actin solution was converted to F-actin by addition of 2 mM MgCl_2_ and 80 mM KCl and incubated overnight at 4 °C. To dissociate tropomyosin, ionic strength was increased to 600 mM KCl and the sample centrifuged at 200,000 × g for 3 h. The resulting pellet was resuspended in G-buffer and extensively dialyzed against G-buffer, then clarified by centrifugation at 400,000 × g for 40 min. G-actin in the final supernatant was re-polymerized by addition of 2 mM MgCl_2_ and 10 mM KCl prior to use.

Actin concentration was determined spectrophotometrically at 290 nm using an extinction coefficient of 26,600 M^−1^·cm^−1^.

### *In vitro* motility assays

To remove Myosin molecules that failed to cycle on actin in the presence of MgATP were removed by pelleting at 350,000 × g for 20 min in the presence of 1.5 mM MgATP and a threefold molar excess of skeletal actin. Using a nitrocellulose-coated flow cell, solutions were applied in 15 μl aliquots in the following sequence. Biotinylated bovine serum albumin (0.5 mg·ml^−1^) in Buffer A (25 mM imidazole pH 7.5, 150 mM KCl, 4 mM MgCl^2^, 1 mM EGTA and 10 mM DTT) was incubated for 1 min, followed by incubation with 5.0 mg·ml^−1^ BSA in Buffer A for 2 min. Neutravidin (25 μg·ml^−1^) in Buffer A was applied for 1 min and then removed by three washes with Buffer A. Myosin (120 μg·ml^−1^ total) in Buffer A was added and incubated for 1 min, then rinsed three times with Buffer A. Rhodamine–phalloidin labeled skeletal muscle actin was applied for 30 s, followed by one wash with Buffer A and one wash with Buffer B (Buffer A supplemented with 0.5% w/v methylcellulose, 25 μg·ml^−1^ PfELC, 25 μg·ml^−1^ PfMLC-B or PfMTIP, 50 μg·ml^−1^ catalase, 125 μg·ml^−1^ glucose oxidase and 3 mg·ml^−1^ glucose).

For standard in vitro motility and motility-mixture experiments, Buffer B containing 2 mM MgATP was introduced twice to initiate motility. For assays at multiple ATP concentrations, Buffer B containing the indicated ATP concentration was added twice to initiate motility. Actin filaments were imaged using an inverted Zeiss Axiovert 10 microscope, a Rolera MGi Plus digital camera, and Nikon NIS-Elements software. Filament velocities were tracked and analyzed with the Fast Automated Spud Trekker (FAST) program (Spudich laboratory, Stanford University; available from http://spudlab.stanford.edu/fast-for-automatic-motilitymeasurements). Velocity distributions were fitted with a Gaussian function.

### Transient kinetics

All kinetic measurements were conducted in buffer containing 10 mM 4-(2-hydroxyethyl)-1-piperazineethanesulfonic acid (HEPES) pH 7.5, 50 mM KCl, 4 mM MgCl_2_, 1 mM EGTA and 1 mM DTT, using a KinTek SF-2004 stopped-flow apparatus (KinTek Corporation). Reported concentrations refer to values after mixing in the stopped-flow cell. Dissociation of actomyosin (0.20 µM actin, 0.15 µM myosin) and actomyosin·ADP (0.20 µM actin, 0.15 µM myosin and 200 µM ADP) was monitored by orthogonal light scattering following rapid addition of 2 mM MgATP. Scattered light was detected at right angles to a 295 nm excitation beam after transmission through a 295 nm interference filter. Mant-nucleotide binding to myosin (0.15 µM) was followed by fluorescence resonance energy transfer (FRET) from a proximal tryptophan residue with excitation at 290 nm and monitoring emission after passing through a 400 nm long-pass filter. Kinetic traces were analyzed with the KinTek software package and fitted to single- or double-exponential functions. Multiple individual time courses (typically 3–8) were averaged prior to curve fitting.

### Crystallization and data collection

To obtain the nucleotide-free (NF) state of the PfMyoB motor domain, 2 mM EGTA was added to the protein solution. Crystals were grown at 4°C using the hanging-drop vapor diffusion method by mixing protein solution (10 mg·ml⁻¹) with crystallization solution at a 1:1 (v:v) ratio. The crystallization solution consisted of 0.1 M HEPES pH 7.5, 1.2 M tri-sodium citrate dihydrate and 8%PEG 400. Prior to data collection, crystals were transferred into a cryoprotectant solution containing 0.1 mM DTT, 0.1 M HEPES pH 7.5, 0.5 M tri-sodium citrate dihydrate, 10% PEG 400, and 25% *glycerol, before flash-freezing in liquid nitrogen*.

X-ray diffraction data were collected at the European Synchrotron Radiation Facility (ESRF) on beamline ID30B^50^ at a wavelength of λ = 0.97625 Å. Data were collected at 100 K using a PILATUS3 6M hybrid photon counting detector (DECTRIS). Diffraction images were processed using the XDS package^51^ and the AutoPROC^52^ pipeline. Crystals of PfMyoB in the nucleotide-free state (PfMyoB-NF) belonged to the orthorhombic space group C222₁. Data collection and refinement statistics are summarized in **Supplementary Table 2**.

### Structure determination and refinement

Molecular replacement was performed with the PfMyoA motor domain in the Rigor-like state (PDB code 6I7D, chain D^9^) without ligand and water as a search model. The solution was modified with SWISS-MODEL^53^ to match PfMyoB sequence (Q8I465). Manual model building and refinement were performed using Coot^54^. Refinement was achieved using phenix refine^55^ from the phenix software suite^56^. The statistics for most favored, allowed and outlier in Ramachandran angles are 97.38%, 2.62% and 0%, respectively.

### All-atom molecular dynamics

The starting point for the all-atom molecular dynamics simulations was based on high-resolution crystal structure of the motor domain of PfMyoB. Two constructs were rebuilt: wild-type motor domain (WT, residues 1-748) and a motor domain with truncated N-terminal extension (ΔN, residues 28-748). CHARMM-gui pipeline (online service) was used to build the all-atom systems which were parameterized with the CHARMM36m forcefield^57,58^. A box consisting of a cube with an edge of 128 Å was used during the simulations. All simulations were performed in the same conditions (150 mM KCl, pH 7.5, with histidine residues described as neutral in the HSD form) and contained PfMyoB motor domain and TIP3 explicit water molecules. Long-range interactions were computed using the particle mesh Ewald (PME) method^59^, facilitating the calculation of the energies.

The simulations were performed under NPT conditions (constant pressure and temperature). The temperature and the pressure of the system were set to 310.15 K with the Nosé-Hoover thermostat and 1 bar with the Parrinello-Rahman barostat^59,60^. The molecular dynamics production timestep was set to 2 fs (dt = 0.002 ps) for all the simulations. The WT and ΔN systems contain 197172 and 197385 atoms respectively. The experiments were performed over 150 ns with GROMACS (version 2023.3)^61^. To ensure the reproducibility of the observations, each simulation was performed in triplicate (see **Supplementary Fig. 5**).

### Trajectory analysis

Trajectories were generated and computed with the module gmx trajconv from GROMACS. Frames of 0.1 ns steps were extracted from the original simulation, that was sampled at 10 ps, for the analysis. All the trajectories were adjusted on the first frame by superimposing on the backbone of the motor domain (28-682) using the macrocommand of VMD (RMSD trajectory tool, version 1.9.4a53, June 29, 2021), allowing the extraction of the RMSD plots. RMSF were obtained with MDanalysis^62^.

All trajectories were analyzed in detail in PyMOL to characterize the evolution of the system and the influence of the N-terminal extension.

### Homology model

Homology model to model the interaction between PfMyoB and KNX-002 was generated using SwissModel^53^, using the structure of PfMyoA complexed to KNX-002 as a template (PDB code 8CDQ^10^).

## Acknowledgement

We are grateful to Sophie Robert, Jean-Louis Rouet and Yohann Brossard from the CaSciModOT service (Calcul Scientifique et Modélisation Orléans-Tours, France) for their technical support and access to the machines. We acknowledge the beamline Id30b from the ESRF (European Synchrotron Radiation Facility) for providing the beamtime to collect diffraction data. We would like to warmly thank Dr. Christoph Mueller-Dieckmann for his assistance during data collection and processing.

## Funding

This work was funded by grants NIH AI132378 (AH and KMT) and ANR (ANR-25-CE44-6310) (JRP).

## Author contributions statement

Conceptualization and design of the research: JRP, KMT, AH. Molecular biology, protein expression and purification: CSB, PMF, JEM. Essential light chain identification, densitometry/stoichiometry and ATPase assays: CSB. Mass spectrometry: MJP. PfMyoB crystallization, data collection and structure determination: DM with the help of AH and JRP. PfMyoB structure refinement: DM, AH, JRP. Functional assays, transient kinetics and *in vitro* motility measurements: JPR. Molecular dynamics simulations: DA.

Formal analysis of the results: JPR, DA, AH, KMT and JRP. The manuscript was written by JRP, KMT, AH and JPR. All the authors reviewed the manuscript. KMT, AH and JRP provided funding. Project administration: KMT, AH and JRP.

## Data availability

The atomic model of PfMyoB in the Rigor state has been deposited in the PDB^63^, under accession code XXX.

## Competing interests

The authors declare no competing interests.

## Notes

### Competing Interest Statement

The authors have declared no competing interest.

