## Supplemental information for "*Plasmodium falciparum* Myosin B is a slow motor optimized for force generation"

### Supplementary information

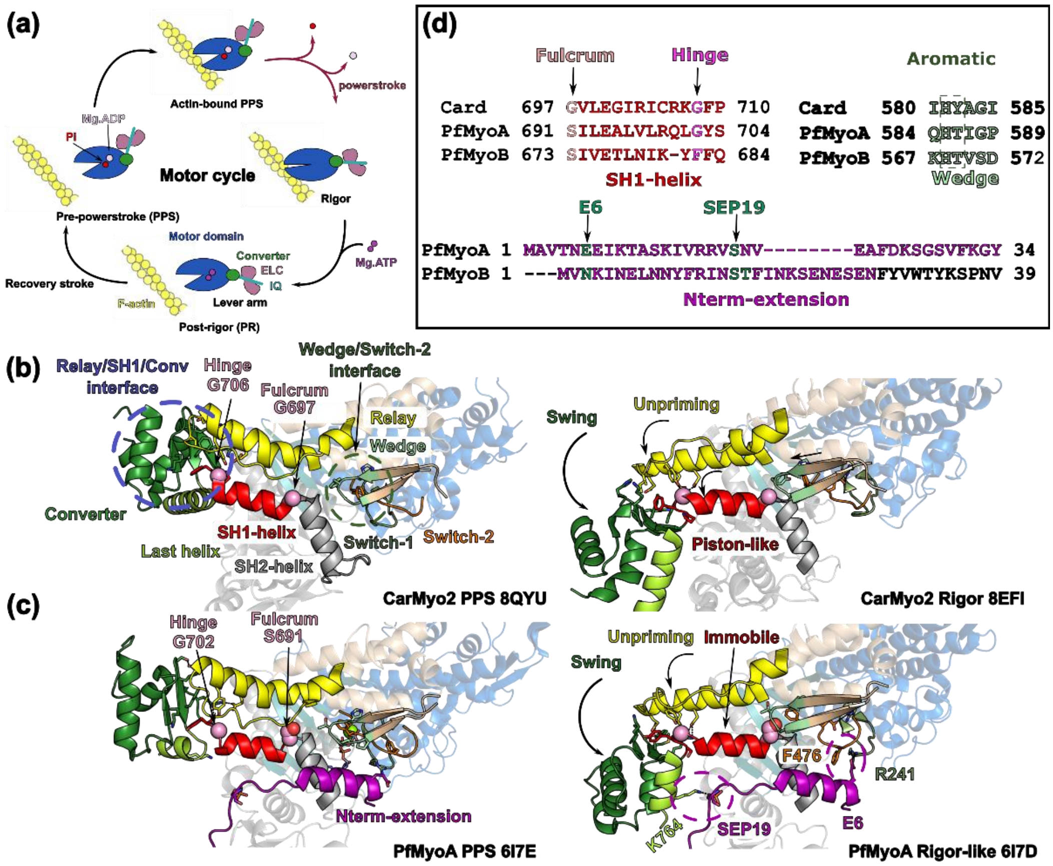

***Supplementary figure 1 – The motor cycle of myosins**. **(a)** Schematic representation of the classical motor cycle of a myosin motors. **(b)** displays the allosteric communication in the conventional cardiac myosin-2 (CardMyo2). In the prepowerstroke state (PPS, PDB code 8QYU[1]), the lever arm, the Relay and the SH1-helix are primed. In the Rigor state (PDB code 8EFI[2]), movements of Switches-1 and -2 from the active site are accompanied by a shift of the Wedge, a piston-like movement of the SH1-helix and an unpriming of the Relay, resulting in the swing of the Converter. The piston-like movement of the lever arm is promoted by the so-called "fulcrum" and the swing of the lever arm is promoted by a hinge, both being flexible glycines. **(c)** In Plasmodium falciparum myosin A (PfMyoA), the glycine of the fulcrum is replaced by a serine, resulting in an immobile SH1-helix. However, the myosin is still able to produce force and perform a swing of the lever arm. This is due to a phoshorylatable N-terminal extension (Nterm-extension) that stabilizes the Rigor state by (i) establishing a polar interaction with K764 from the Converter and (ii) E6 binds R241, that is involved in a cation-π interaction with F476, stabilizing the Switches. PfMyoA PPS and Rigor-like states (respectively PDB codes 6I7E and 6I7D[3]) are illustrated in **(c)**. (d) shows sequence divergences in the allosteric connectors (SH1-helix and Wedge) in PfMyoA and human β-cardiac myosin (Card). Plasmodium falciparum myosin B (PfMyoB), another class-XIV myosin is also shown. PfMyoB also shows divergences in the connectors, but also differs from PfMyoA, especially in the sequence of the Nterm-extension, where two residues could be phosphorylated (S16 and T17).*

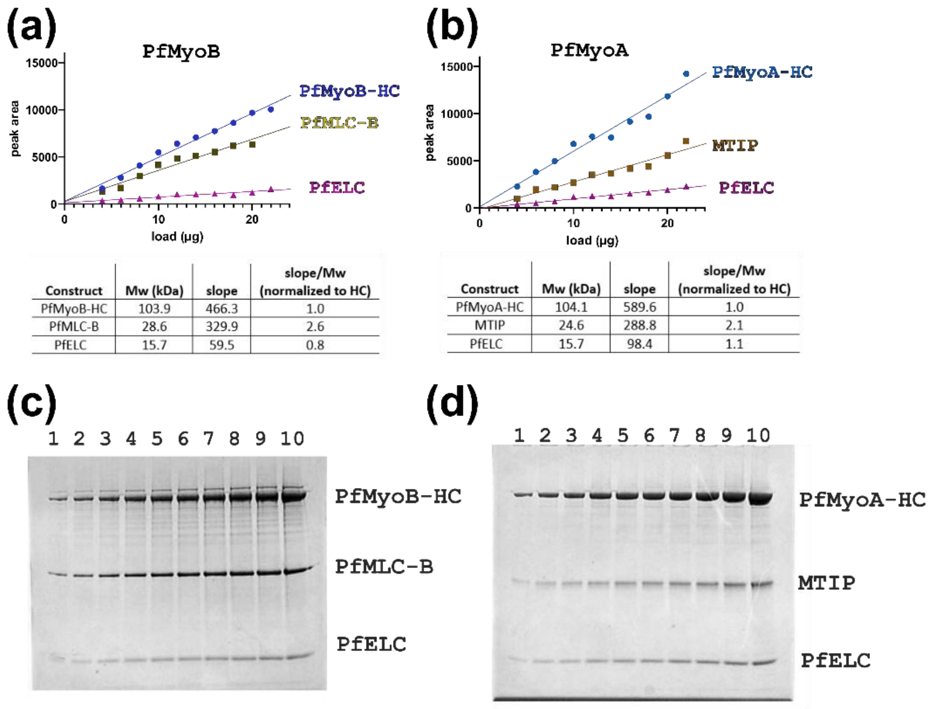

| Construct | Mw (kDa) | slope | slope/Mw (normalized to HC) |
|---|---|---|---|
| PfMyoB-HC | 103.9 | 466.3 | 1.0 |
| PfMLC-B | 28.6 | 329.9 | 2.6 |
| PfELC | 15.7 | 59.5 | 0.8 |

| Construct | Mw (kDa) | slope | slope/Mw (normalized to HC) |
|---|---|---|---|
| PfMyoA-HC | 104.1 | 589.6 | 1.0 |
| MTIP | 24.6 | 288.8 | 2.1 |
| PfELC | 15.7 | 98.4 | 1.1 |

***Supplementary figure 2 – Gel densitometry analysis of PfMyoB-HC/MLC-B/PfELC and PfMyoA-HC/MTIP/PfELC.*** ***(a)*** *and* ***(b)*** *Gel densitometry analysis of SDS-PAGE gels of PfMyoB-HC/MLC-B/PfELC* ***(a)*** *and PfMyoA-HC/MTIP/PfELC* ***(b)****.* ***(c)*** *and* ***(d)*** *Coomassie stained 12% SDS-PAGE gels of PfMyoB/PfMLC-B/PfELC* ***(c)*** *and PfMyoA/MTIP/PfELC* ***(d)****. Total protein loaded was 4 ug (lane 1), 6 µg (lane 2), 8 µg (lane 3), 10 µg (lane 4) 12 µg (lane 5), 14 µg (lane 6), 16 µg (lane 7), 18 µg (lane 8), 20 µg (lane 9) and 22 µg (lane 10).*

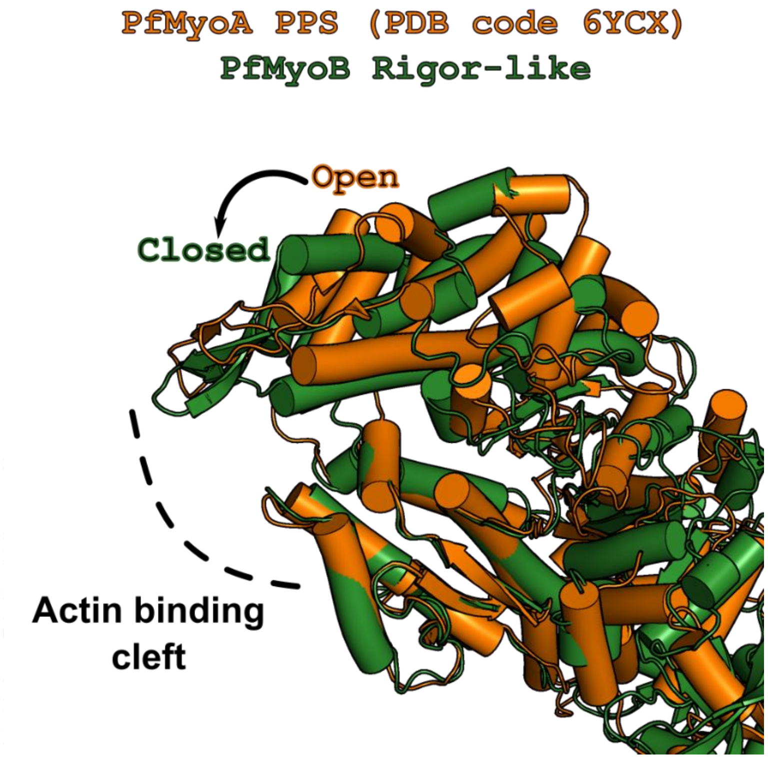

***Supplementary figure 3 – The crystal structure of PfMyoB in the nucleotide-free condition has the actin binding cleft close.*** *The structure of the PPS of PfMyoA with actin binding cleft open and the structure of PfMyoB in the nucleotide-free condition have been superimposed on the L50 subdomain. The superimposition shows that PfMyoB has the actin binding cleft closed, matching all the criteria of a Rigor-like state.*

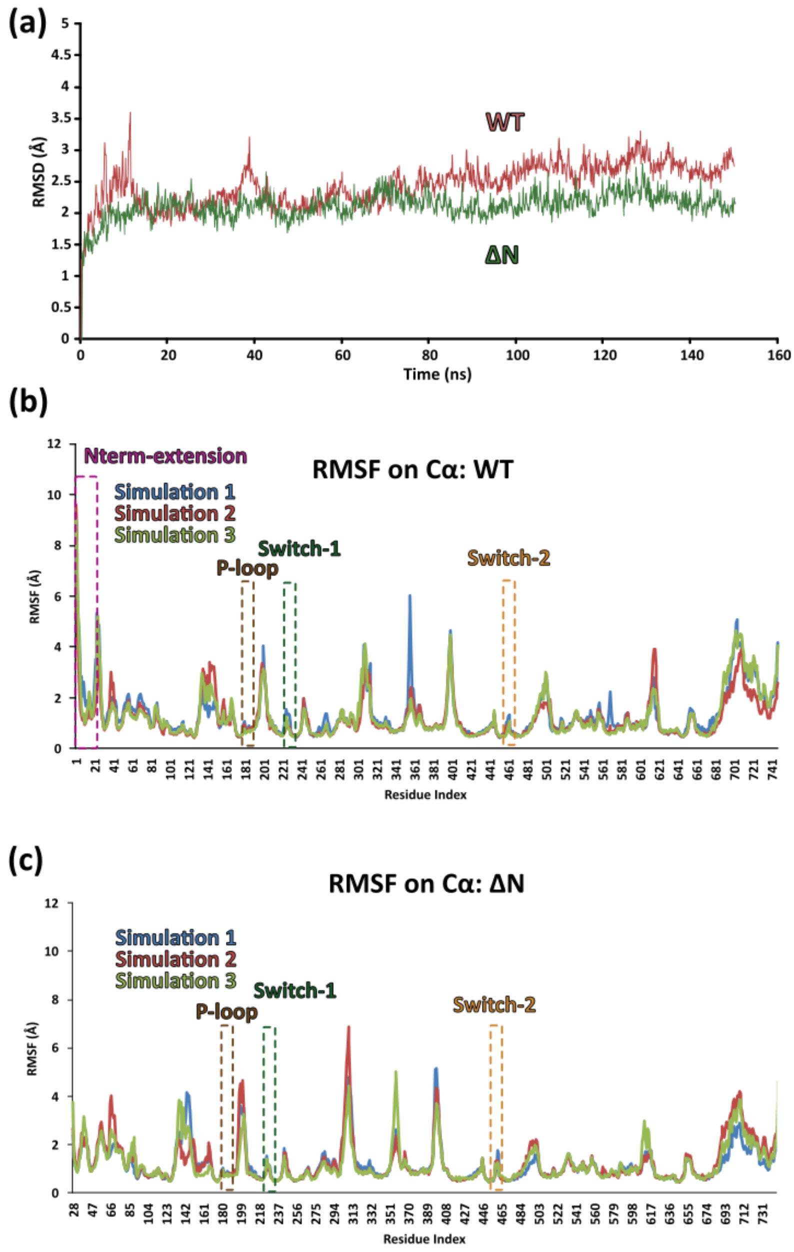

***Supplementary figure 4 – All-atom molecular dynamics simulation of the PfMyoB motor domain. (a)*** *shows the root mean square deviation (RMSF) calculated on the Cα for the two constructs. In each condition, two frames (t1 and t2) were selected to illustrate how the N-terminal extension restrains the conformational exploration of the three connectors of the active site: P-loop, Switch-1 (Sw1) and Switch-2 (Sw2). The root mean square fluctuations (RMSF) were computed on Cα from 150 ns trajectories in two conditions to evaluate the effect of the N-terminal extension:* ***(b)*** *the wild-type motor domain (WT) and* ***(c)*** *the motor domain without the N-terminal extension (ΔN). Experiments were reproduced three times (simulations 1-3) to ensure the reproducibility of the results. On each graph, the positions of the three active site connectors are indicated (P-loop, Switch-1 and Switch-2), the position of the N-terminal extension (Nterm-extension) is indicated.*

| Name | Gene Identifier from PlasmoDB | Function | Reference (name in reference) |
|---|---|---|---|
| Pf CAM | PF3D7_1434200 | Calmodulin-like | Pires et al, 2022[4] (PLC7) |
| PfCAM-ΔAll | Modified from PF3D7_1434200 | Calmodulin-like | Pires et al, 2022[4] (PLC7) with E31Q, E67Q, E104Q, E140Q mutations |
| LC-1320c | PF3D7_0627200 | Unknown | Pires et al, 2022[4] (PLC5) |
| Pf ELC | PF3D7_1017500 | PfMyoA essential light chain | Bookwalter et al, 2017[5], Robert-Paganin et al, 2019[3], Moussaoui et al, 2023[6] |
| Pf LC-2 | PF3D7_0816400 | Unknown | |
| Pf LC-3 | PF3D7_0728500 | Unknown | |
| Pf LC-4 | PF3D7_0605400 | calcium binding protein, putative/troponin C | Hall et al, 2002[7] |
| Pf LC-5 | PF3D7_0414200.2 | Calmodulin-like | Pires et al, 2022[4] (PLC3) |
| Pf LC-6 | PF3D7_0714400 | Calmodulin-like | |
| Pf LC-7 | PF3D7_1030800 | Calmodulin-like | |

***Supplementary Table 1** – List of putative essential light chains used in co-expression with PfMyoB and PfMLC-B.*

| | PfMyoB-NF |
|---|---|
| **Data collection** | |
| Space group | $C222_1$ |
| Cell dimensions | |
| *a*, *b*, *c* (Å) | 136.568, 262.681, 83.619 |
| α, β, γ (°) | 90.00, 90.00, 90.00 |
| Resolution (Å) | 121.170 - 2.040 (2.233 – 2.040) |
| $R_{merge}$ (all I+ & I-) | 0.138 (1.968) |
| $R_{merge}$ (within I+ & I-) | 0.135 (1.887) |
| Number of observations (total) | 867553 (43241) |
| Number of observations (unique) | 68398 (3421) |
| $\langle I / \sigma I \rangle$ | 12.4 (1.6) |
| Completeness (Spherical) (%) | 71.4 (15.2) |
| Completeness (Ellipsoidal) (%) | 93.8 (58.7) |
| Redundancy | 12.7 (12.6) |
| $CC_{1/2}$ | 0.997 (0.555) |
| **Refinement** | |
| Resolution (Å) | 70.54 - 2.04 (2.113 – 2.04) |
| No. reflections | 68397 (337) |
| $R_{work}$ / $R_{free}$ | 0.1738/0.2011 |
| No. atoms | 6446 |
| Protein | 6126 |
| Ligand/ion | 21 |
| Water | 299 |
| *B*-factors (Å$^2$) | 61.04 |
| Protein | 61.46 |
| Ligand/ion | 60.25 |
| Water | 52.50 |
| R.m.s. deviations | |
| Bond lengths (Å) | 0.008 |
| Bond angles (°) | 0.94 |

**Supplementary Table 2. Data collection, phasing and refinement statistics for PfMyoB-NF.** Data were collected at the ESRF synchrotron, on the ID30b beamline[8].

| | Association rate constant ($\mu M^{-1} s^{-1}$) | Dissociation rate constant ($s^{-1}$) | Affinity ($\mu M$) |
|---|---|---|---|
| | | MgATP | |
| *MyoA WT* | 5.3 ± 0.1 | 0.6 ± 0.4 | 0.1 ± 0.1 |
| ***MyoB WT*** | 3.7 ± 0.1 | 2.0 ± 0.4 | 0.5 ± 0.1 |

| | Association rate constant ($\mu M^{-1} s^{-1}$) | Dissociation rate constant ($s^{-1}$) | Affinity ($\mu M$) |
|---|---|---|---|
| | | MgADP | |
| *MyoA WT* | 4.9 ± 0.1 | 4.0 ± 0.8 | 0.8 ± 0.2 |
| ***MyoB WT*** | 3.5 ± 0.1 | 4.7 ± 0.3 | 1.3 ± 0.1 |

***Supplementary table 3 – Association and dissociation rate constants for mant-ATP (top) and mant-ADP binding (bottom) to PfMyoB WT and PfMyoA WT.*** *Temperature, 30°C. Values, mean ± SE of the fit. PfMyoA WT data from*[6].
